# TNF-α induces type I IFN signalling to suppress neurogenesis and recruit T cells

**DOI:** 10.1101/2025.06.18.660440

**Authors:** Tinne Amalie Damgaard Nissen, Arishma Baig, Daniel T. Rock, Sandra Shibu, Hyunah Lee, Lauren A. O’Neill, Susan John, Linda S. Klavinskis, Sandrine Thuret

## Abstract

Adult hippocampal neurogenesis (AHN) is essential for learning, memory, and mood regulation, and its disruption is implicated in ageing, neurodegeneration, and mood disorders. However, the mechanisms linking inflammation to AHN impairment remain unclear. Here, we identify chronic tumour necrosis factor-alpha (TNF-⍺) signalling as a key driver of neurogenic dysregulation via a previously unrecognised type I interferon (IFN) autocrine/paracrine loop in human hippocampal progenitor cells (HPCs). Using a human *in vitro* neurogenesis model, single-cell RNA sequencing, and functional T cell migration assays, we show that TNF-⍺ induces a robust type I IFN response in HPCs, promoting chemokine-mediated, and CXCR3-dependent T cell recruitment and suppressing neurogenesis. This inflammatory signalling cascade drives a fate switch in HPCs from a neurogenic trajectory towards an immune-defensive phenotype, with critical implications for infectious and inflammatory disease pathogenesis. These findings uncover a key inflammatory checkpoint regulating human AHN and highlight potential therapeutic targets to restore neurogenesis in chronic inflammatory states.

## Introduction

Adult neurogenesis occurs in two main neurogenic niches in the mammalian brain: the subventricular zone (SVZ) lining the lateral ventricles and the subgranular zone (SGZ) of the dentate gyrus of the hippocampus^1^. Upon activation, hippocampal neural stem cells (NSCs) known as radial glia-like cells (RGLs) divide and differentiate into astrocytes or excitatory, granule cells, integrating into the granule cell layer (GCL). Initially discovered in rodents^2^, human adult hippocampal neurogenesis (AHN) has since been demonstrated using carbon dating^3^, postmortem tissue^4–7^, *in vitro* propagation of adult neural progenitor cells (NPCs)^8^, single nucleus RNA sequencing^9^, and spatial transcriptomics^10^. AHN in rodents enhances several cognitive functions, including memory and learning^11^, the development of individuality^12^, and mood regulation^13^.

Perturbations in AHN observed in ageing, neurodegenerative diseases, and mood disorders are thought to contribute to cognitive decline and associated symptoms, given AHN’s role in memory and mood regulation^14, 15^. Experimental enhancement of AHN reduces depressive-like behaviours^16^ and rescues memory and emotional deficits in Alzheimer’s disease (AD) mice^17^. Emerging evidence suggests that inflammation, elevated in AD^18^, ageing^19^, and in some individuals with mood disorders such as depression^20^, may underlie these AHN deficits. In particular, the proinflammatory cytokine TNF-⍺ has been implicated in both AD^21, 22^ and depression^23, 24^, and is known to negatively regulate AHN^25, 26^. These observations support a model in which inflammation-driven suppression of AHN contributes to shared clinical phenotypes across disorders. Targeting this axis may represent a viable neuroprotective strategy^15^. Defining the molecular mechanisms by which inflammatory mediators, such as TNF-⍺, influence hippocampal neural stem cell (NSC) fate is crucial for developing interventions preserving AHN and its associated cognitive functions in inflammatory conditions.

Chronic TNF-⍺-mediated inflammation was recently shown to drive a functional switch in rodent olfactory NSCs, redirecting them from neuroregeneration toward immune recruitment^27^. These NSCs actively recruited inflammatory cells, including T cells, at the expense of their neurogenic capacity. We hypothesised that a similar switch could occur in human hippocampal progenitor cells (HPCs), contributing to impaired AHN and T cell infiltration observed in ageing and neurodegenerative disease, where TNF-⍺ levels are persistently elevated^28–30^. Using an established *in vitro* model of human hippocampal neurogenesis^31–36^, we found that chronic TNF-⍺ exposure suppressed neurogenesis and induced robust CXCL10 secretion. Single-cell RNA (scRNA) sequencing revealed that TNF-⍺ activated type I IFN signalling via autocrine/paracrine signalling, a mechanism not previously described in central nervous system (CNS)-resident cells. This signalling cascade drove the secretion of chemokines that mediated CXCR3-dependent T cell chemotaxis and contributed to the anti-neurogenic effects of chronic TNF-⍺. Together, our findings support a model in which chronic TNF-⍺ reprograms human HPCs from a neurogenic to a proinflammatory state, orchestrated by TNF-⍺-induced type I IFN signalling.

## Results

### TNF-⍺-driven functional and phenotypic reprogramming of HPCs

To investigate how TNF-⍺ reprograms human HPCs, we first confirmed that our cellular model expressed the TNF-⍺ receptors TNFR1 and TNFR2 (Supplementary Fig. 1). TNF-α stimulation of HPCs induced rapid nuclear translocation of NF-κB p65, peaking at 30 minutes and remaining elevated for up to 48 hours (Fig. 1a–b), consistent with sustained activation of a canonical proinflammatory pathway implicated in the regulation of neurogenesis^37^. To assess the functional relevance of TNF-⍺ signalling in HPCs and their progeny, we evaluated the secretion of the NF-κB-regulated cytokine IL-6 and a panel of NF-κB-regulated chemokines. Levels were measured in the supernatant at 24 and 48 hours after TNF-⍺ treatment of the HPCs, as well as 24 hours, 48 hours, and seven days of TNF-⍺ treatment in differentiating HPCs (Supplementary Fig. 2). Among the chemokines analysed (Supplementary Fig. 3-5), CXCL10 exhibited the most robust induction in response to TNF-⍺ stimulation (Fig. 1c–d). To examine the effects of chronic TNF-⍺ on neuronal differentiation, HPCs were pre-treated ± TNF-⍺ under proliferative conditions for 48 hours, followed by seven days of differentiation with continued TNF-⍺ exposure every 48 hours (Fig. 1e). TNF-⍺ dose-dependently reduced the proportion of doublecortin (DCX)+ neuroblasts (Fig. 1g), while only the highest dose (10 ng/ml) decreased the percentage of cells expressing the neuronal marker microtubule-associated protein 2 (MAP2) (Fig. 1h). In parallel, TNF-⍺ reduced total neurite length in both populations, indicating impaired morphological maturation (Supplementary Fig. 6). Collectively, these findings suggest that chronic TNF-⍺ functionally and phenotypically reprograms HPCs.

**Fig. 1.**
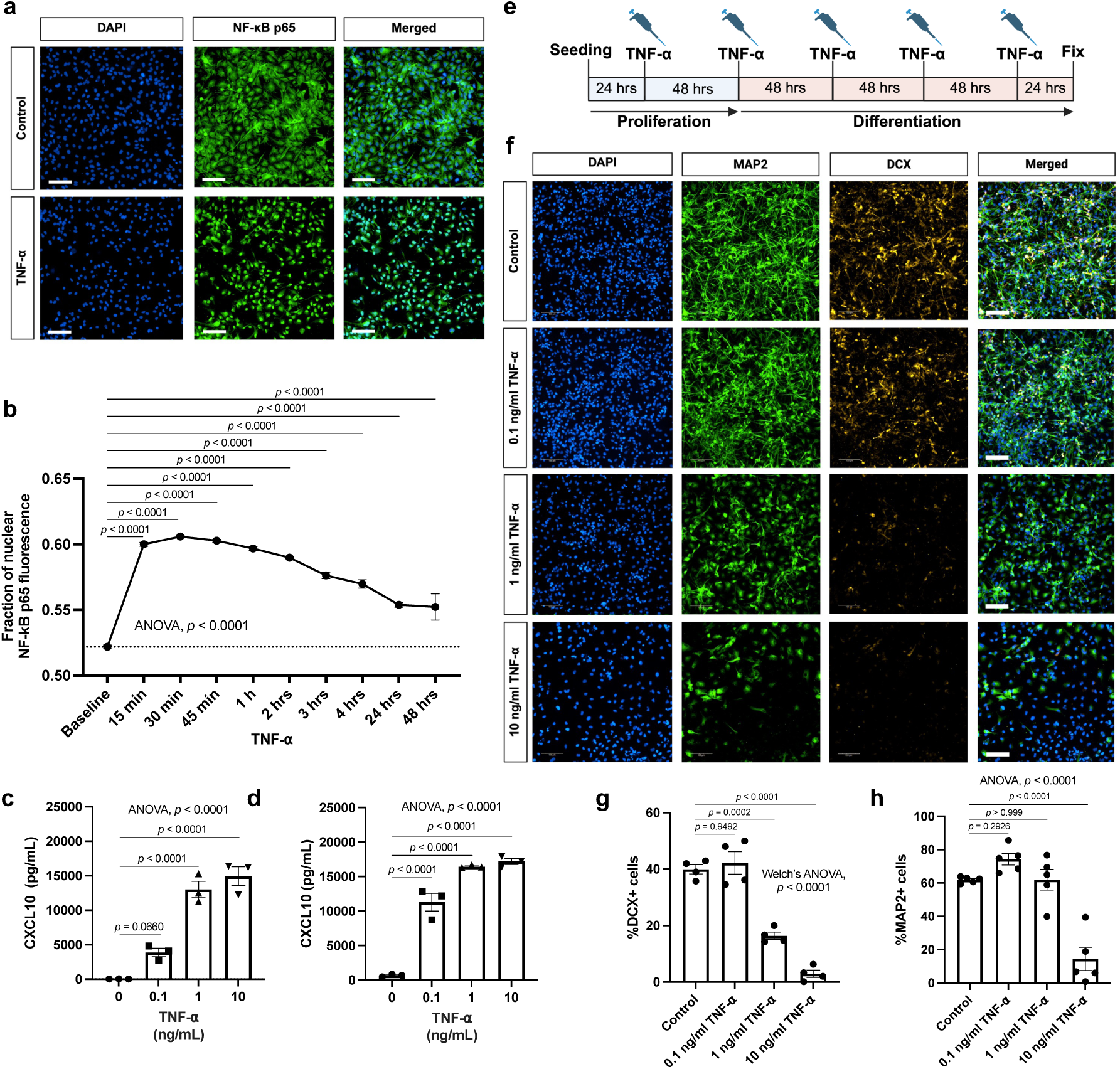
TNF-⍺ drives immune signalling and decreases neurogenesis in human hippocampal progenitor cells. **a**, Representative immunocytochemistry images of three independent experiments with similar results showing the expression of NF-κB p65 on HPCs treated ± 10 ng/ml TNF-⍺ for 15 minutes. Scale bar represents 100 µm. **b**, Quantification of the fraction of nuclear NF-κB p65 fluorescence intensity relative to total cellular fluorescence. Data represent mean ± SEM from *n* = 3. Statistical analysis: one-way ANOVA followed by Bonferroni multiple comparisons test. **c**, and **d**, Quantification of CXCL10 levels in culture supernatants measured by ELISA. **c**, HPCs under proliferation conditions treated for 48 hours ± 0.1, 1, or 10 ng/ml TNF-⍺. **d**, HPCs under differentiation conditions treated for 7 days ± 0.1, 1, or 10 ng/ml TNF-⍺. Data represent mean ± SEM from *n* = 3. Statistical analysis: one-way ANOVA followed by Bonferroni multiple comparisons test. **e**, Schematic overview of the chronic TNF-⍺ treatment regime for investigating its effect on neuronal differentiation. HPCs were treated under proliferation conditions for 48 hours ± 0.1, 1, or 10 ng/ml TNF-⍺ before inducing differentiation for seven days under treatment ± 0.1, 1, or 10 ng/ml TNF-⍺ every 48 hours. **f**, Representative images showing the expression of MAP2 (green) and DCX (orange) on seven days differentiated HPCs treated chronically ± 0.1, 1, or 10 ng/ml TNF-⍺. Scale bar represents 100 µm. **g**, and **h**, Quantification of the percentage of DCX+ cells and MAP2+ cells respectively based on **f**. Data represent mean ± SEM from. **g**, Data based on *n* = 4. Statistical analysis: Welch’s ANOVA followed by Games-Howell post-hoc test for pairwise comparisons. **h**, Data based on *n* = 5. Statistical analysis: one-way ANOVA followed by Bonferroni multiple comparisons test.

### Chronic TNF-α disrupts neuronal differentiation and upregulates a transcriptional type I IFN signature

To elucidate the molecular mechanisms underlying the effects of chronic TNF-⍺, we performed scRNA sequencing of HPCs exposed for 48 hours to low (0.1 ng/ml) or high (1 ng/ml) TNF-⍺, as well as their progeny after 7 and 14 days of differentiation under chronic TNF-⍺ exposure (Fig. 2a). After quality control, 22,133 cells were identified across 15 clusters, including astrocyte-like (clusters 1-4), radial glial-like (RGL)/NSC-like (5-6), intermediate progenitor cell (IPC)-like (7-10), neuroblast-like (11), and immature neuron-like (12) clusters (Fig. 2b-d). The neurogenic identity of clusters 11 and 12 was confirmed by overlap with a human immature granule cell signature^9^ and by enrichment of neuronal biological processes and cellular compartment gene ontology (GO) terms (Fig. 2e-g and Supplementary Fig. 7). Critically, HPCs differentiated in the presence of high-dose TNF-⍺ showed a marked reduction in cells within neurogenic clusters (Fig. 2h), demonstrating disrupted neurogenesis.

**Fig. 2.**
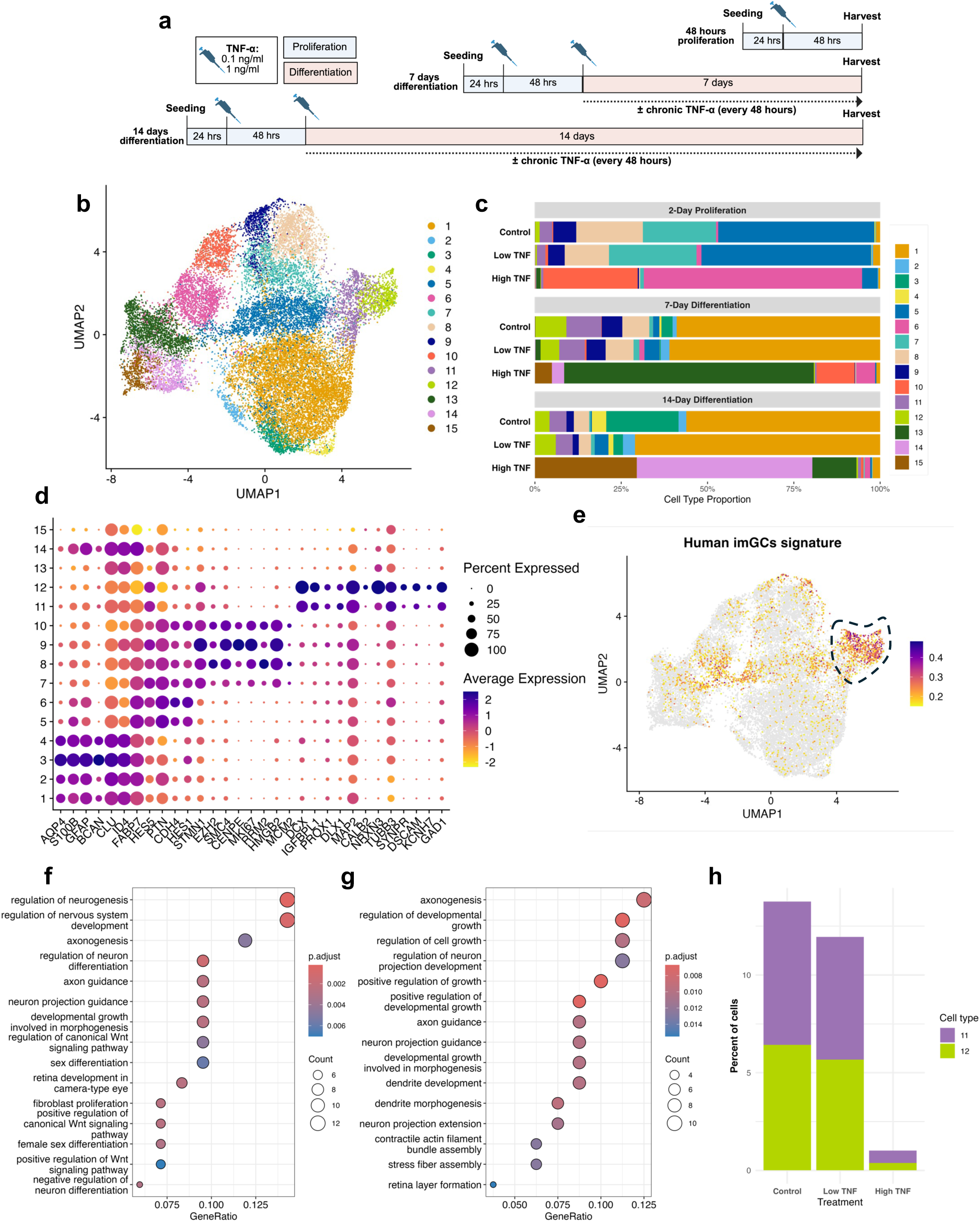
Chronic TNF-⍺ impairs neuronal differentiation. **a**, Schematic overview of the experimental design. **b**, UMAP plot showing transcriptional clustering of single cells. Colors indicate the 15 cell clusters identified. **c**, Stacked bar graphs showing the frequencies of cell clusters shown in **b** across the time and treatment groups. **d**, Dot plot showing the expression of canonical marker genes for expected cell types across the cell clusters. **e**, UMAP plot showing each cell scored against a human immature granule cells (imGCs) signature^9^. The outlined area highlight the of the signature with cluster 11 and 12. **f**, and **g**, dot plots showing GO enrichment analysis of biological process terms based on the top 100 genes positively defining cluster 11 and cluster 12 respectively. **h**, Bar plots show the percentage cells belonging to the neurogenic cluster 11 (green) and 12 (purple) within differentiation samples from 7 and 14 days by treatment condition: control, low TNF (0.1 ng/ml), and high TNF (1 ng/ml).

While control and low-dose TNF-⍺ treated cells clustered together, high-dose TNF-⍺ treated cells formed distinct clusters (Extended Fig. 1a). RGL-like cluster 6 and IPC-like cluster 10, primarily composed of HPCs treated with high-dose TNF-⍺, exhibited strong upregulation of type I interferon (IFN) response genes including *ISG15*, *IFI27*, and *IFI6* (Fig. 3a-b). GO enrichment analysis highlighting “response to type I IFNs” and phagocytic vesicle components (Extended Fig. 1c-d; Supplementary Fig. 8). The type I IFN signature was detected across clusters specific to treatment with high-dose TNF-⍺ (Fig. 3c). HPCs differentiated with high-dose TNF-⍺ failed to align with astrocyte-, neuroblast-, or immature neuron-like clusters, indicating disrupted lineage commitment and transcriptional dysregulation. Go term analysis of clusters 13-15 indicated a shift toward aberrant, immune-reactive states with activation of a type I IFN-mediated antiviral program (Fig. 3d-f). Given that TNF-⍺ does not directly induce type I IFN response genes, we hypothesised that a type I IFN autocrine/paracrine loop mediated the ISG upregulation. HPCs and their progeny expressed the machinery to respond to type I IFNs (Supplementary Fig. 9), and cells from high-dose TNF-⍺-specific clusters strongly overlapped with an NSC-specific IFN-β response signature^38^ (Fig. 3g). Transcription factor activity analysis predicted STAT1, STAT2, and IRF9 activity in these clusters, consistent with IFN signalling (Fig. 3h). Together, these findings demonstrate that chronic TNF-⍺ exposure induces type I IFN response genes in HPCs, suppresses neurogenesis, and promotes aberrant immune-reactive cellular states during differentiation.

**Fig. 3.**
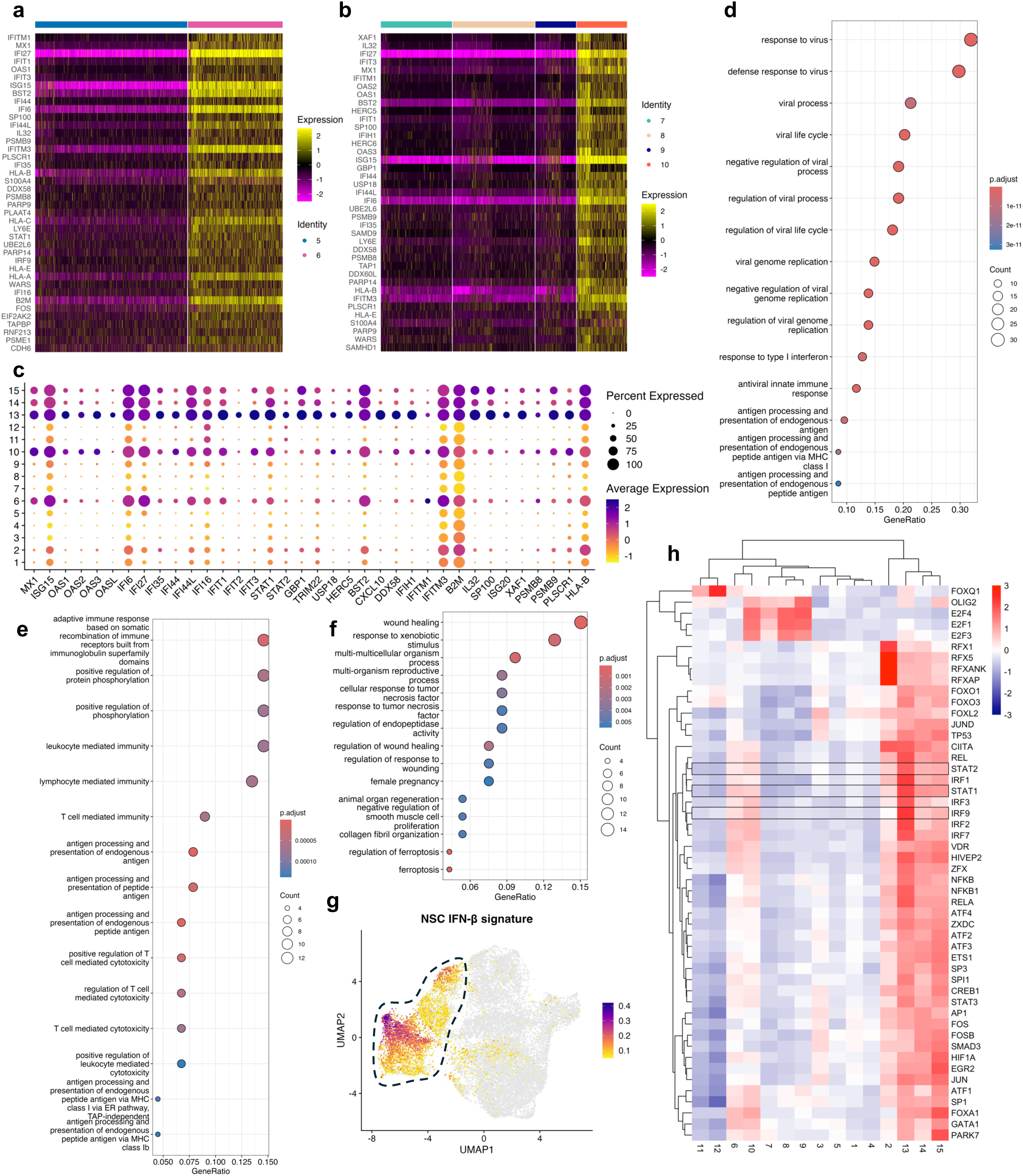
Chronic TNF-⍺ drives the upregulation of type I IFN genes. **a**, Heatmap of top 40 upregulated genes based on log fold change in cluster six as compared to cluster five. **b**, Heatmap of top 40 upregulated genes based on log fold change in cluster ten as compared to cluster seven, eight, and nine. **c**, Dot plot showing the expression of interferon response genes across the cell clusters. **d**, Dot plot showing gene ontology (GO) enrichment analysis of biological processes terms based on the top 100 genes positively defining cluster 13. **e**, Dot plot showing gene ontology (GO) enrichment analysis of biological processes terms based on the top 100 genes positively defining cluster 14. **f**, Dot plot showing gene ontology (GO) enrichment analysis of biological processes terms based on the top 100 genes positively defining cluster 15. **g**, UMAP plot showing each cell scored against a NSC IFN-β transcriptional signature. The outlined area highlight the of the signature with cluster 6, 10, 13, 14, and 15. **h**, Heatmap showing the mean activity per cluster of the top 50 more variable transcription factors. STAT1, STAT2 and IRF9 is highlighted.

### TNF-⍺ drives type I IFN signalling through activation of IFNAR

As STAT1 negatively regulates human hippocampal neurogenesis^39^ and STAT1/STAT2 regulates the expression of type I IFN-regulated genes, we assessed their activation in HPCs following TNFa stimulation. Phosphorylation of both STAT1 and STAT2 was delayed, peaking at 3–6 hours (STAT1: Fig. 4a–b and Supplementary Fig. 11; STAT2: Extended Fig. 2a–b and Supplementary Fig. 12; GAPDH stability in response to TNF-⍺: Supplementrary Fig. 10), consistent with a mechanism of indirect activation. In line with an autocrine/paracrine loop, factors secreted downstream of TNF-⍺-mediated receptor engagement increased the percentage of ISG15+ HPCs (Extended Fig. 2c–d). TNF-⍺ transiently induced *IFNB1* expression, peaking at 3 hours (Fig. 4b), temporally correlating with STAT1/STAT2 activation, suggesting IFN-β as the likely mediator. TNF-⍺-induced *IFNB1* expression depended on NF-κB signalling (Extended Fig. 2e) but was independent of *de novo* protein synthesis (Extended Fig. 2f). As IRF1 is required for TNF-α–induced *IFNB1* expression in other cell types^40, 41^, we examined its regulation in HPCs. TNF-α markedly upregulated *IRF1*, peaking at 3 hours (Supplementary Fig. 13), preceding *IFNB1* induction and suggesting a potential regulatory role for IRF1 in HPC *IFNB1* induction. TNF-⍺-induced upregulation of the STAT1/STAT2-regulated chemokine CXCL10 has been found to depend on IFN-β^42^. Consistent with this, the delayed induction of *CXCL10* in response to TNF-α in HPCs required *de novo* protein synthesis, as shown by its sensitivity to cycloheximide treatment (Fig. 4d-e). To confirm that TNF-⍺ induces type I IFN signalling via an autocrine/paracrine loop, we blocked the IFN-⍺/β receptor (IFNAR) using the antibody anifrolumab^43^. Anifrolumab fully abolished TNF-⍺-induced STAT1+ cell induction and significantly reduced ISG15+ cell upregulation (Fig. 4f-h). Similarly, Janus kinase (JAK) inhibition downstream of IFNAR using JAK Inhibitor I^44^ blocked TNF-⍺-induced ISG15+ HPC upregulation (Extended Fig. 2g-h). Finally, TNF-⍺ upregulated the type I IFN-induced surface marker tetherin/BST-2 on HPCs (Extended Fig. 2i and Supplementary Fig. 14), which was recently shown to be elevated on dysfunctional NSCs in ageing neurogenic niches^28^. Collectively, these findings demonstrated that TNF-⍺ drives autocrine/paracrine type I IFN signalling in HPCs via IFNAR and downstream JAKs likely mediated by IFN-β.

**Fig. 4.**
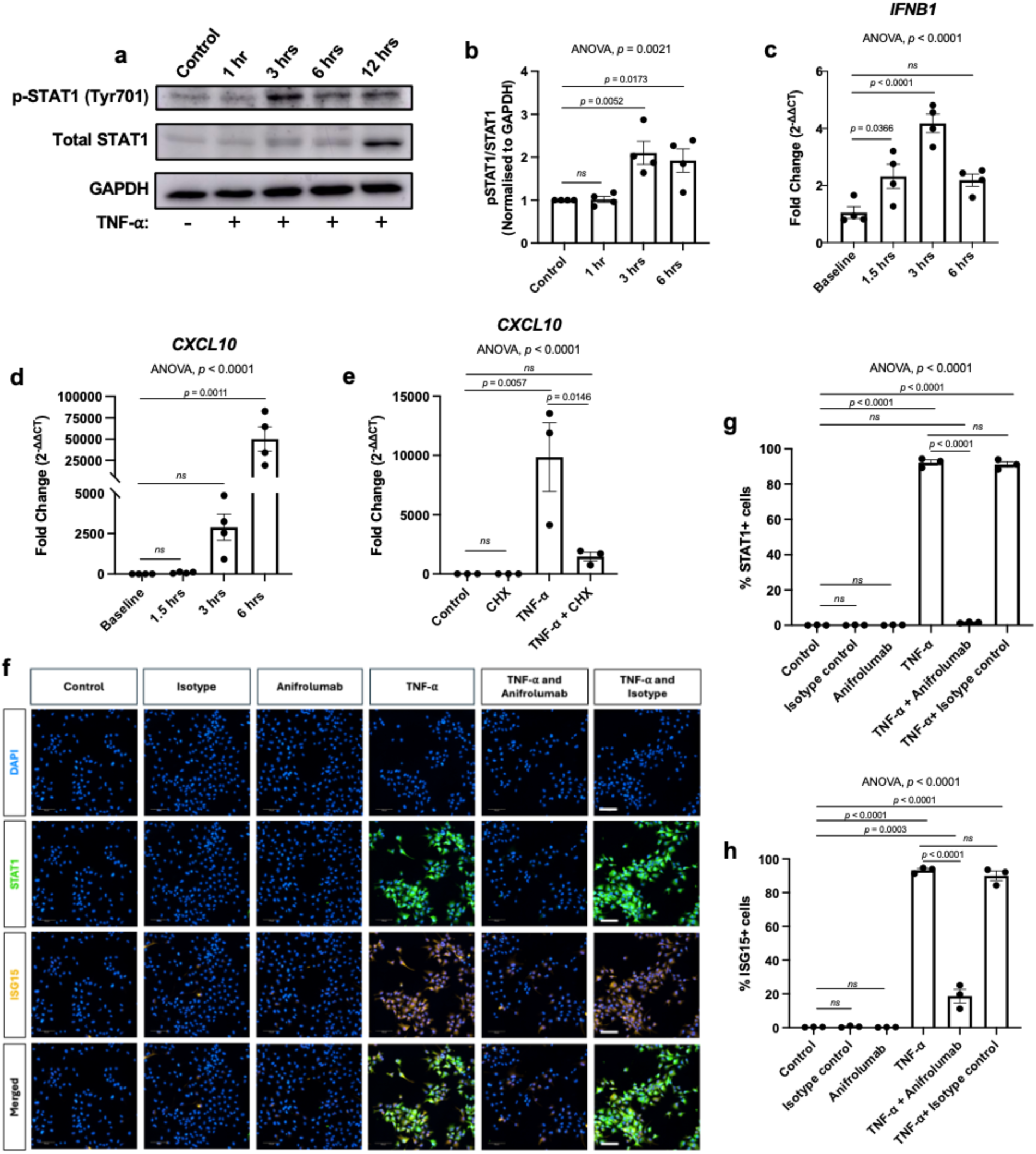
TNF-⍺ drives type I IFN signalling in human hippocampal progenitor cells via autocrine/paracrine signalling via IFNAR. **a**, Representative western blots of four independent experiments showing the activation of STAT1 in HPCs treated with 1 ng/ml TNF-⍺ for 1 hr, 3 hrs, 6 hrs, and 12 hrs. Cell lysates were probed for phosphorylated STAT1 (p-STAT1 Tyr701), total STAT1, and GAPDH as a loading control. **b**, Quantification of STAT1 activation in human hippocampal progenitor cells from **a**. pSTAT1 and STAT1 intensities were normalised to GAPDH intensity before calculating the fold change pSTAT1/STAT1 relative to control. Data represent mean ± SEM from four independent experiments. Statistical analysis: one-way ANOVA followed by Bonferroni’s multiple comparisons. **c**, RT-qPCR data showing the relative gene expression of *IFNB1* in response to treatment with 1 ng/ml TNF-⍺. Data is represented as mean ± SEM of four independent experiments. Statistical analysis: one-way ANOVA followed by Bonferroni’s multiple comparisons. **d**, RT-qPCR data showing the relative gene expression of *CXCL10* in response to treatment with 1 ng/ml TNF-⍺. Data is represented as mean ± SEM of four independent experiments. Statistical analysis: Kruskal-Wallis test followed by Dunn’s post hoc correction. **e**, RT-qPCR data showing the relative gene expression of *CXCL10* in HPCs treated for six hours with 10 µg/ml CHX or DMSO ± 1 ng/ml TNF-⍺. Data is represented as mean ± SEM of three independent experiments. Statistical analysis: one-way ANOVA followed by Bonferroni’s multiple comparisons. **f**, Representative images of three independent experiments with similar results showing the expression of STAT1 (green) and ISG15 (orange) on HPCs. Scale bar represents 100 µm. **g**, Quantification of the percentage of STAT1+ cells based on **f**. Data represent mean ± SEM. Statistical analysis: one-way ANOVA followed by Bonferroni’s multiple comparisons. **h**, Quantification of the percentage of ISG15+ cells based on **f**. Data represent mean ± SEM. Statistical analysis: one-way ANOVA followed by Bonferroni’s multiple comparisons.

### TNF-⍺-conditioned media from HPCs and differentiated progeny drive CXCR3-dependent T cell chemotaxis

After elucidating the molecular mechanism of TNF-⍺-induced type I IFN signalling in HPCs, we next investigated its functional consequences. Recent studies highlight T cell infiltration into neurogenic niches in ageing and AD, with T cells found near NSCs^28, 30, 45–49^. We found high levels of the type I IFN-regulated chemokines CXCL10 and CXCL11 in the supernatant of HPCs and their differentiated progeny following TNF-⍺ treatment. As both chemokines promote T cell chemotaxis via the CXCR3 receptor ^50^, we hypothesised that their secretion could mediate CXCR3-dependent T cell recruitment. We established an *in vitro* transwell assay using primary human T cells from three healthy donors and validated that recombinant CXCL10 promotes CXCR3-dependent chemotaxis of both CD4+ and CD8+ T cells (Supplementary Fig.15-16).

We generated conditioned media (CM) from TNF-⍺–treated HPCs as well as from HPC-derived cells differentiated under chronic TNF-⍺ exposure, and assessed its ability to induce CXCR3-dependent migration of activated primary human CD4+ and CD8+ T cells (Fig. 5a–h). CM from both TNF-⍺–treated HPCs and their differentiated progeny significantly enhanced the chemotactic index of CD4+ and CD8+ T cells. This effect was significantly blunted by pre-treatment of the T cells with a CXCR3 antagonist, indicating that the migratory response was primarily CXCR3-dependent. These findings suggest that both HPCs and their differentiated progeny secrete chemokines capable of promoting T cell migration through CXCR3 signaling.

**Fig. 5.**
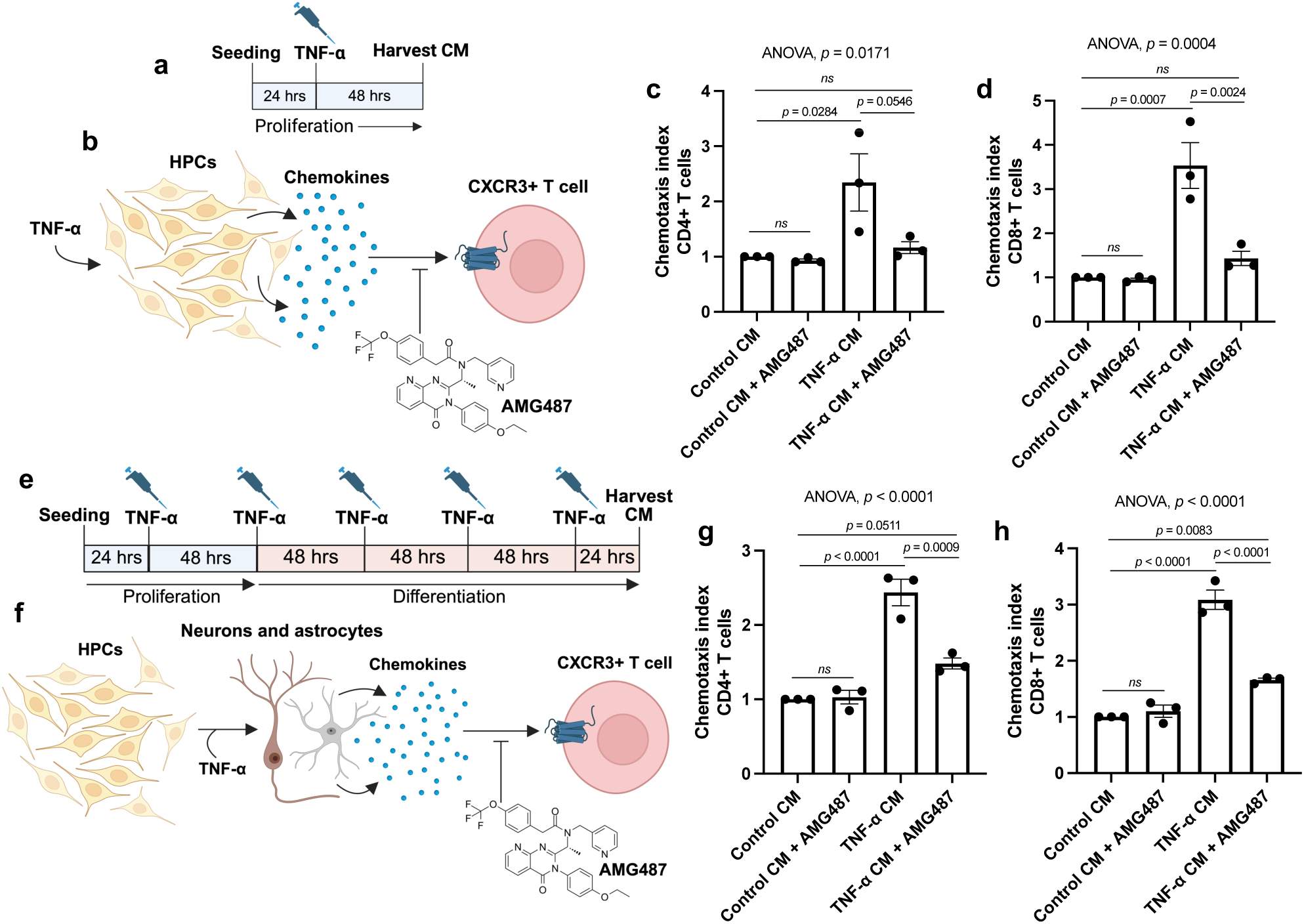
TNF-⍺-conditioned media from proliferating and differentiating HPCs drive CXCR3-dependent chemotaxis of primary, human T cells. **a**, Schematic overview of the experimental design for generating conditioned media (CM) from HPCs treated for ± 1 ng/ml TNF-⍺ for 48 hrs. **b**, Chemokine secretion by HPCs in response to TNF-⍺ stimulation assessed for its ability to induce CXCR3-dependent T cell chemotaxis. **c**, and **d**, Quantification of the chemotactic response (chemotaxis index) of **c** CD4+ T cells and **d** CD8+ T cells treated ± 1 μM AMG487 or DMSO in response to control CM or TNF-⍺-CM from HPCs. Data is represented as mean ± SEM of three healthy donors. Statistical analysis: one-way ANOVA followed by Bonferroni’s multiple comparisons test. **e**, Schematic overview of the experimental design for generating CM from differentiated HPCs treated chronically ± 1 ng/ml TNF-⍺. **f**, Chemokine secretion by differentiated HPCs in response to chronic TNF-⍺ stimulation assessed for its ability to induce CXCR3-dependent T cell chemotaxis. **g**, and **h**, Quantification of the chemotactic response (chemotaxis index) of **g** CD4+ T cells and **h** CD8+ T cells treated ± 1 μM AMG487 or DMSO in response to control CM or TNF-⍺-CM from differentiated HPCs. Data is represented as mean ± SEM of three healthy donors. Statistical analysis: one-way ANOVA followed by Bonferroni’s multiple comparisons test.

As our data indicated TNF-⍺-induced *CXCL10* upregulation was mediated via the type I IFN autocrine/paracrine signalling loop, we tested whether IFNAR or JAK inhibition abrogated T cell chemotaxis in response to TNF-⍺-CM from HPCs (Fig. 6a-b). Pharmacological blockage of both IFNAR and JAK independently abrogated the chemotactic response of CD4+ T and CD8+ T cells towards chemokines secreted by TNF-⍺-stimulated HPCs (Fig. 6c-d). These results indicated that the secretion of CXCR3 ligands by HPCs in response to TNF-⍺ was dependent on the autocrine/paracrine type I IFN signalling loop.

**Fig. 6.**
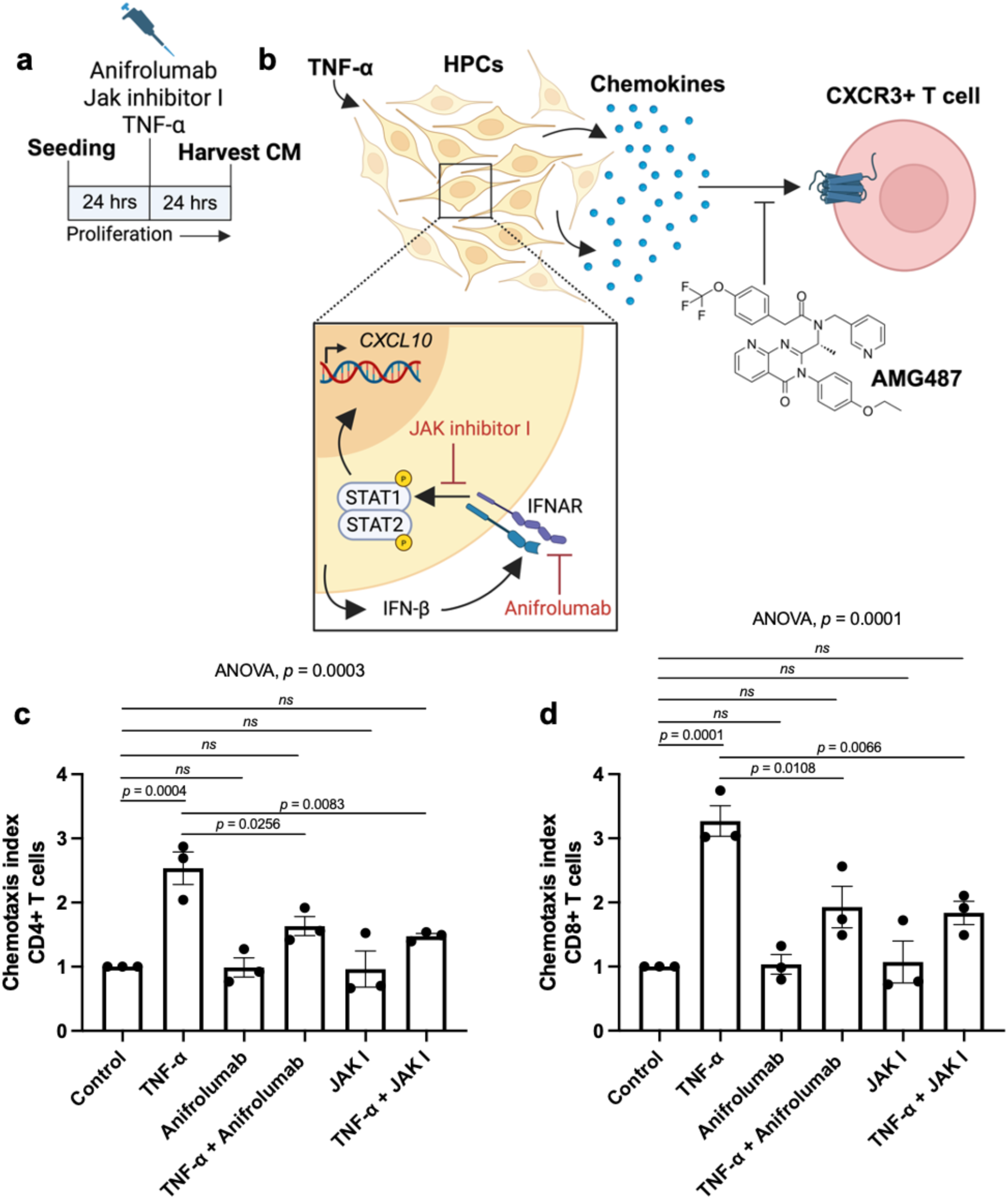
JAK and IFNAR signalling are indispensable for TNF-⍺-induced HPC chemokine secretion directing T cell chemotaxis. **a**, Schematic overview of the experimental design for generating conditioned media (CM) from HPCs treated ± 1 ng/ml TNF-⍺, ± 10 µg/ml anifrolumab (IFNAR antagonist), and ± 0.16 nM JAK inhibitor I for 24 hrs. **b**, The effect of IFNAR and JAK inhibition was evaluated for its ability to abrogate CXCR3-dependent T cell chemotaxis towards HPC TNF-⍺-CM. **c**, and **d**, Quantification of the chemotactic response (chemotaxis index) of **c** CD4+ T cells and **d** CD8+ T cells in response to the CM generated as outlined in **a**. Data is represented as mean ± SEM of three healthy donors. Statistical analysis: one-way ANOVA followed by Bonferroni’s multiple comparisons test.

In addition to chemokine secretion, TNF-⍺ upregulated the expression of adhesion molecules in HPCs implicated in T cell interactions including *ICAM1* and *VCAM1* (Supplementary Fig. 17). Notably, *ICAM1* expression was significantly increased after 3-6 hours of TNF-⍺ exposure and remained elevated on the HPC surface at 24 hours post-treatment (Supplementary Fig. 18b).

Together, these findings demonstrated that TNF-⍺ stimulation of HPCs promoted CXCR3-dependent chemotaxis of primary human T cells *in vitro* through a type I IFN-mediated autocrine/paracrine loop. Furthermore, TNF-⍺ enhanced HPC expression of adhesion molecules, potentially facilitating direct HPC–T cell interactions.

### Blocking IFNAR signalling rescues DCX⁺ neuroblast loss induced by chronic TNF-⍺

Given the prominent activation of STAT1/STAT2 downstream of TNF-⍺, and the established role of STAT1 as a negative regulator of hippocampal neurogenesis and cognition^39, 51–53^, we hypothesised that chronic TNF-⍺ may exert its antineurogenic effects via IFNAR-dependent type I interferon signalling. To test this, we administered the IFNAR-blocking antibody anifrolumab, which effectively suppressed TNF-⍺–induced ISG15 expression in differentiating HPCs (Supplementary Fig. 19), confirming inhibition of type I IFN signalling. As previously observed, chronic exposure to 1 ng/ml TNF-⍺ selectively impaired the generation of DCX⁺ neuroblasts without affecting the proportion of MAP2⁺ neurons (Fig. 7a–c). Notably, IFNAR blockade fully rescued the loss of DCX⁺ neuroblasts, implicating autocrine/paracrine type I IFN signalling as a key mediator of TNF-⍺–driven neurogenic deficits. These findings are consistent with prior evidence that disrupting type I IFN pathways can restore hippocampal neurogenesis in ageing models^54^.

**Fig. 7.**
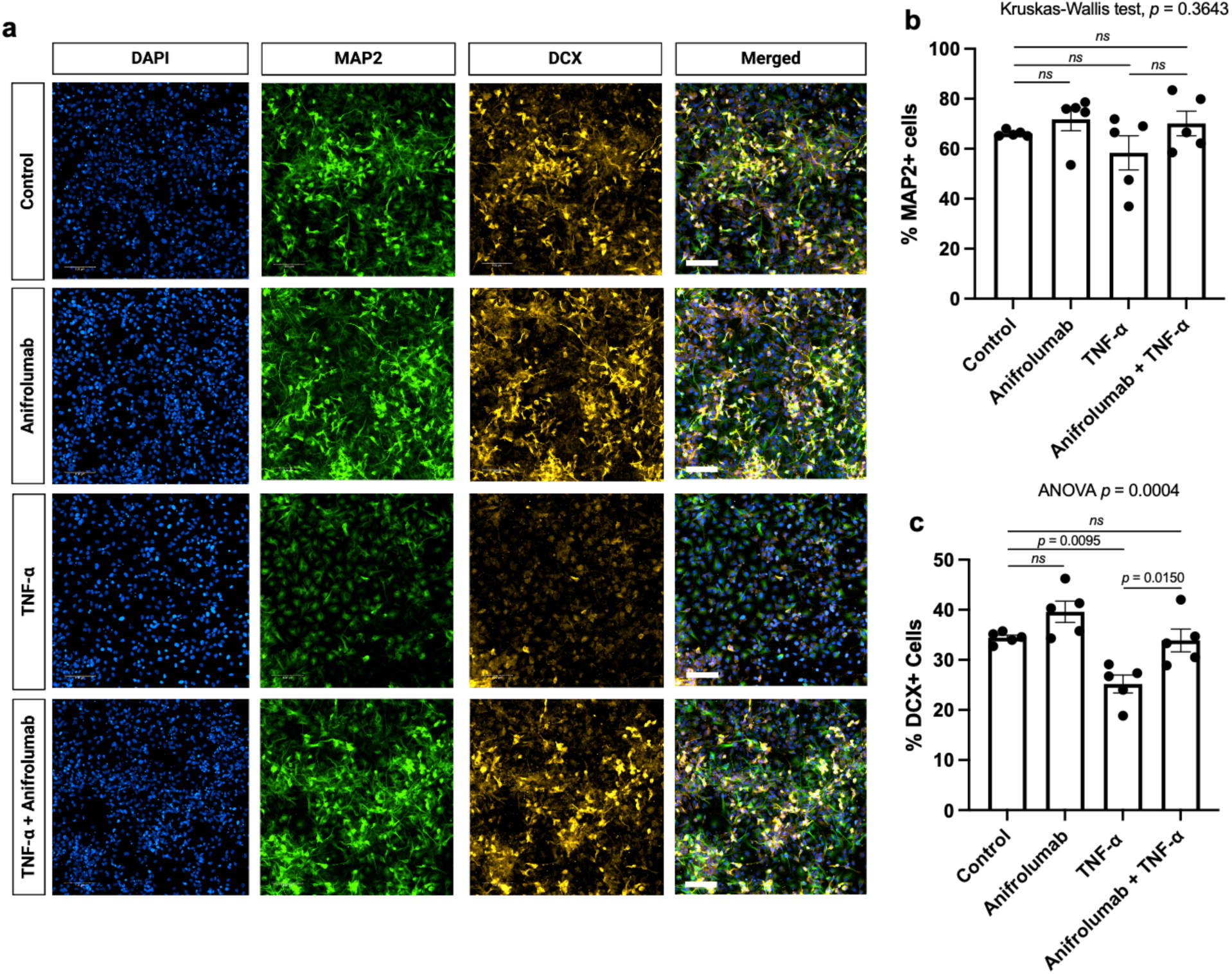
IFNAR blockage rescues the chronic TNF-⍺-induced decrease in the percentage of DCX+ neuroblasts. **a**, Representative images showing the expression of MAP2 (green) and DCX (orange) ion seven days differentiated HPCs treated chronically ± 1 TNF-⍺ ± 10 µg/ml anifrolumab. Scale bar represents 100 µm. **b**, and **c,** Quantification of the percentage of **b** MAP2+ cells and **c** DCX+ cells respectively based on **a**. Data represent mean ± SEM from five independent experiments. Statistical analysis: **b** Kruskal-Wallis test followed by Dunn’s post hoc correction and **c** one-way ANOVA followed by Bonferroni’s multiple comparisons test.

## Discussion

Emerging evidence implicates neuroinflammation, impaired AHN, and T cell accumulation within the hippocampal neurogenic niche as key features of ageing and neurodegenerative diseases^28–30^. However, the mechanistic links connecting these phenomena remain unclear. Here, we hypothesised that chronic TNF-⍺ exposure functionally reprogrammes HPCs, diverting them from a neurogenic fate toward an immune-defensive phenotype, parallelling recent observations in olfactory NSCs^27^. This phenotypic switch may both impair AHN directly and promote T accumulation, which could further suppress neurogenesis. Together, these effects may contribute to cognitive deficits observed in ageing and neurodegeneration. Utilising a human *in vitro* model we demonstrated that chronic TNF-⍺ suppress hippocampal neurogenesis while promoting CXCL10 secretion. To investigate the molecular mechanisms underlying these observations, we performed scRNA sequencing, which unexpectedly revealed a type I IFN transcriptional signature in HPCs and their differentiating progeny. Mechanistically, we found that TNF-⍺ engages a type I IFN signalling cascade in HPCs, driven by an autocrine/paracrine IFN-β. Moreover, this pathway suppressed neurogenesis and induced CXCR3 ligand secretion, promoting chemotaxis of activated primary human T cells. Our data uncovered a previously uncharacterised TNF-⍺-IFN-β-IFNAR-CXCL10 axis that may exist in other CNS-resident cells and its role in reprogramming HPCs from a neurogenic to an immune-defensive state under chronic inflammatory conditions. Importantly, IFNAR antagonism restored TNF-⍺-induced neuroblasts loss and blocked T cell recruitment, identifying IFNAR signalling as a dual-action candidate therapeutic target for mitigating inflammation-induced neurogenic dysfunction.

The antineurogenic effect of TNF-⍺ we observed closely mirrors findings from primary NSCs and *in vivo* models^26, 55–57^, reinforcing the validity of our human cellular system. Similarly, the TNF-⍺-induced impairment of neuronal morphological maturation^57, 58^ and CXCL10 secretion align with previous reports in primary cells^59^. These parallels underscore the physiological relevance of our model and support its utility for mechanistic dissection of inflammation-driven neurogenic dysfunction. Future *in vivo* studies will be critical to extend these findings.

While type I IFN signalling is well characterised in infectious diseases^60^, its role in sterile inflammation remains less understood, but increasingly recognised^61, 62^. Our findings identify a TNF-⍺-IFN-β-IFNAR axis in human HPCs, representing a novel mechanism of type I IFN activation by an endogenous inflammatory cytokine in a CNS-resident progenitor cell. Similar TNF-⍺-induced IFN-β responses have been reported in non-CNS cell types, requiring IRF1 and NF-κB ^40–42, 63^. Consistent with this, we showed that TNF-⍺ induces *IFNB1* expression in HPCs via an NF-κB–dependent, protein synthesis– independent mechanism that may involve IRF1. Whether the TNF-⍺-IFN-β-IFNAR axis operates in CNS cell types beyond HPCs—such as microglia, where IFN signatures have recently been identified across a broad range of sterile brain pathologies^64^ remains an important question for future investigations. Given IFNAR’s ubiquitous expression^65^, HPC-derived IFN-β may act on surrounding niche cells, potentially amplifying local inflammatory crosstalk.

Although type I IFNs are traditionally considered as antiviral and neuroprotective, sustained or dysregulated signalling has been increasingly implicated in ageing^54, 66^ and AD^67–69^. Elevated type I IFNs impair AHN^39, 54^ and promote cognitive deficits^54, 68, 70^ both of which can be reversed through IFNAR antagonism^54, 68^. These cognitive impairments are likely mediated, at least in part, by IFN-induced AHN dysfunction. Furthermore, type I IFNs can directly promote amyloidogenic, neurotoxic processes^71^. Our data support the pathogenic role for type I IFN signalling: notably, we found that TNF-⍺ suppresses neurogenesis via type I IFN signalling and shifts differentiating progenitors towards an IFN-response, astroglial-like state. This is consistent with recent evidence that Zika virus, which drives type I IFN signalling, promotes a molecular switch from neurogenesis towards astrogliogenesis^72^. Thus, type I IFNs may represent a converging downstream mechanism through which chronic inflammation impairs AHN and contributes to cognitive decline in ageing and neurodegeneration.

T cells infiltrate neurogenic niches in the context of ageing and AD^28–30, 45–49, 73^, and have been demonstrated to modulate brain ageing and neuroregeneration^74, 75^. Understanding the mechanisms underlying their infiltration and interactions with brain resident cells is therefore critical. We found that TNF-⍺ treatment induces HPC chemokine secretion promoting CXCR3-dependent chemotaxis of activated primary human T cells, suggesting hippocampal NSCs may directly recruit T cells during neuroinflammation, consistent with findings in murine olfactory NSCs^27^. This indicates a conserved mechanism across neurogenic niches and species, where chronic TNF-⍺ reprograms NSCs towards immune cell recruitment. In addition to NSCs, microglial activation is reported to drive CXCL10-mediated CD8+ T cell recruitment and promote aging-related white matter degeneration^76^. Thus, multiple CNS cell types likely cooperate in recruiting T cells via CXCR3 ligands in ageing and neurodegeneration.

We demonstrated that IFNAR and JAK antagonism abolished T cell chemotaxis towards TNF-⍺-stimulated HPC-derived CM, demonstrating that autocrine/paracrine type I IFN signalling is the primary driver of CXCR3 ligand secretion. This mechanism parallels findings in a nephritis model, where TNFR2-induced IFN-β autocrine signalling promotes renal monocyte recruitment^41^. Of note, in the brain, type I IFN signalling in astrocytes promotes brain metastasis by enhancing monocytic myeloid cell recruitment^77^. Beyond promoting T cell chemotaxis, HPC-secreted CXCR3 ligands may exert direct deleterious effects on within the neurogenic niche, as CXCR3 activation has been demonstrated to drive hippocampal neural hyperexcitability^69, 78, 79^.

As CXCL10 and CXCL11 are upregulated by IFN-γ^50^, IFN-γ-producing CD8+ T cells infiltrating ageing neurogenic niches^28^ may promote further CXCR3 ligand secretion from NSCs, potentially creating a feed-forward loop enhancing T cell recruitment. This may explain why CD8+ T cells have been found in close proximity to NSCs^28^. Increasing the number of CD8+ T cells in the hippocampus has been shown to decrease the number of proliferative progenitors and DCX+ newborn neurons^29^. This effect is likely mediated by IFN-γ secretion which has been shown to negatively affect NSCs^28^. Consistently, CXCL10 inhibition enhances neuroblast production^80^, possibly by reducing T cell infiltration. Thus, TNF-⍺ may impair AHN both by driving antineurogenic type I IFN signalling in NSCs and recruiting CD8+ T cells that suppress neurogenesis via IFN-γ. Blocking adaptive immune cell infiltration into neurogenic niches may represent a therapeutic strategy for cognitive decline in ageing and neurodegeneration. Given the low abundance of NSCs, future *in vivo* studies are needed to determine their contribution to T cell recruitment in neuroinflammation.

The TNF-⍺-induced functional switch we observed in human HPCs —from a neurogenic to an immune responsive phenotype —may represent a protective adaptation to infection or injury. Since neurogenesis is energetically costly, redirecting cellular resources to mount an immune response could be advantageous for resolving the underlying pro-inflammatory challenge.

Collectively, our results provide novel insights into how chronic TNF-⍺ changes the fate of human HPCs from neurogenesis towards immune functions, elucidating the molecular mechanism underlying this switch. Furthermore, this work offers a novel explanation for the emerging appearance of type I IFN signalling in ageing and neurodegeneration where TNF-⍺ levels are elevated and highlights HPCs as potential direct participants in the progression of chronic inflammation.

## Methods

### Cellular model of hippocampal neurogenesis

We used the human hippocampal progenitor cell line (HPC0A07/03A; ReNeuron LTD., Surrey, United Kingdom), derived from a first-trimester female foetus ethically approved under UK and US guidelines, as a cellular model for hippocampal neurogenesis as previously described^33, 34, 36, 81^. The cell line was immortalised using the c*-mycER^TAM^* gene to conditionally proliferate in presence of 100 nM 4-hydroxy-tamoxifen (4-OHT) (Sigma Aldrich, Cat No. H7904), 10 ng/mL human basic fibroblast growth factor (bFGF) (Peprotech, Cat No. 100-18B-500) and 20 ng/mL human epidermal growth factor (EGF) (Peprotech, Cat No. AF100-15–500) and spontaneously differentiate in their absence. The cells were cultured as previously described^82^ on laminin (Gibco, Cat No. 23017015) coated culture vessels (Nunclon, Denmark) at 37 °C, 5 % CO2.

### TNF-⍺-treatments

The HPCs were treated with TNF-ɑ (Peprotech, Cat No. 300-01A) for the duration and with the concentrations indicated in the individual experiments.

### Immunocytochemistry

Cells were washed with PBS and fixed in 4% paraformaldehyde (in PBS) for 15 min at room temperature. Cells were permeabilised and blocked for 1 h at room temperature in blocking buffer: 0.3% Triton X-100 (Sigma, Cat No. X100) and 5% normal donkey serum (Bio-Rad, Cat No. C06SB) in PBS. For TNFR1 and TNFR2 staining, a non-permeabilising block (5% normal donkey serum in PBS) was used. Primary antibodies (Supplementary Table 1), diluted in blocking buffer, were applied overnight at 4 °C. The following day, cells were washed 3× with PBS and incubated with secondary antibodies (Supplementary Table 1) in blocking buffer for 2 h at room temperature, protected from light. 300 nM DAPI (Sigma, Cat No. 62248) in PBS was applied for 5 min at room temperature. Cells were then washed 4× with PBS and stored at 4 °C in PBS containing 0.05% sodium azide (Merck, Cat No. S2002).

### High-content imaging and analysis

Immunostained cells were imaged using the Opera Phenix Plus High-Content Screening System (PerkinElmer). Imaging was performed in a single focal plane using a 20× water-immersion objective. A minimum of 9 fields per well were acquired from 96-well plates and averaged to generate a mean per well. Channels were captured using the following excitation/emission settings: DAPI (ex 385 nm/em 425–475 nm), Alexa Fluor 488 (ex 488 nm/em 500–550 nm), and Alexa Fluor 555 (ex 561 nm/em 609–644 nm). A minimum of three technical replicates per condition were imaged and averaged for each biological replicate. Images were analysed using Harmony software (PerkinElmer). Briefly, cells were segmented using the *Find Nuclei* building block. Morphological parameters were calculated using *Calculate Morphological Properties*, and nuclei smaller than 50 μm² or larger than 500 μm² were excluded to remove apoptotic bodies and artefacts. Cells at image borders were also excluded. Cytoplasmic or nuclear signal intensity was quantified using *Find Cytoplasm*, *Find Morphological Properties*, and *Calculate Intensity Properties* building blocks. Positive cells were identified using the *Select Population* building block by applying a per-plate intensity threshold based on control wells. The percentage of positive cells was calculated using the *Define Results* building block.

### RNA extraction, cDNA synthesis, and RT-qPCR

HPCs (3.5 × 10⁵ cells per well) were seeded in 6-well plates coated with 20 μg/mL mouse laminin (Sigma, Cat No. L2020,). Cells were treated the following day as indicated. For lysis, media were removed, wells were washed with PBS, and cells were lysed in 350 μL Buffer RLT Plus (Qiagen, Cat No. 74134) supplemented with 1:100 2-mercaptoethanol (Bio-Rad, Cat No. 1610710). Lysates were scraped, transferred to microcentrifuge tubes, vortexed, and stored at –80 °C. Total RNA was extracted using the RNeasy Plus Mini Kit (Qiagen, Cat No. 74134,) following the manufacturer’s instructions, and RNA concentrations were measured using a NanoDrop spectrophotometer (Thermo Fisher Scientific). Extracted RNA (200 ng) was reverse transcribed using the High-Capacity cDNA Reverse Transcription Kit (Applied Biosystems, Cat No. 4368813) following the manufacturer’s protocol. The thermal cycler program (Biometra Trio series, Analytik Jena) used for cDNA synthesis is provided in Supplementary Table 2. qPCR reactions contained 5 μL cDNA (1:4 dilution in nuclease-free water), 10 μL SYBR Green Master Mix (PCR Biosystems, Cat No. PB20.11-05,), 4 μL nuclease-free water, and 1 μL primer mix (10 pmol), in a total volume of 20 μL. Reactions were run in triplicate in 384-well plates (Applied Biosystems, Cat No. 4309849,) on a QuantStudio 5 Real-Time PCR System (Applied Biosystems) using the cycling protocol described in Supplementary Table 3. Data were analysed with QuantStudio Design & Analysis Software (v1.5.1), and relative gene expression was calculated using the 2^–ΔΔCt method with GAPDH as the reference gene. Primer sequences are provided in Supplementary Table 4.

### Western blotting

HPCs (3–6 × 10⁵ cells per well) were seeded in laminin-coated 6-well plates (Thermo Fisher Scientific, Cat No. 140675). Cells were treated the following day as indicated. Cells were washed with PBS and lysed in 150 μL 1 × Laemmli buffer (Bio-Rad, Cat. No. 1610747) diluted in PBS and supplemented with 355 mM β-mercaptoethanol (Bio-Rad, Cat. No. 1610710). Lysates were scraped, boiled for 5 min at 95 °C, and stored at –80 °C until analysis. Proteins were resolved by 10% SDS–PAGE and transferred to PVDF membranes (Thermo Fisher Scientific, Cat. No. 88518). Membranes were blocked for 1 h at room temperature in PBS containing 5% milk and 0.1% Tween-20, then washed 3× (10 min each) with PBS + 0.5% Tween-20. Primary antibodies (Supplementary Table 5) were applied overnight at 4 °C in blocking buffer. After washing, membranes were incubated with HRP-conjugated secondary antibodies for 1 h at room temperature, followed by 3× washes in PBS + 0.5% Tween-20. Detection was performed using Clarity Max ECL (Bio-Rad, Cat. No. 1705062,) for STAT proteins and Clarity ECL (Bio-Rad, Cat. No. 1705061) for GAPDH, each for 4 min. Membranes were imaged on an ImageQuant 800 Imager (Amersham), and band intensities were quantified using Image Studio.

### Western blotting quantifications

Band intensities for phosphorylated and total STAT proteins, as well as GAPDH, were measured using Image Studio. Phosphorylated and total STAT signals were normalised to the corresponding GAPDH signal. The ratio of phosphorylated to total STAT protein was then calculated for each sample and expressed relative to control.

### Statistical analysis

Analyses were performed using GraphPad Prism (v10.4.1) and R (v4.3.2) with rstatix (v0.7.2). Normality was assessed by Q–Q plots and Shapiro–Wilk test, and variance homogeneity by Levene’s test. Results are reported as mean ± SEM, with statistical significance set at p < 0.05. The specific test and sample size are stated in each figure legend. For normally distributed data, unpaired Student’s t-test was used for two-group comparisons, and one-way ANOVA with Bonferroni post hoc for >2 groups. If variances were unequal, we applied Welch’s t-test or Welch’s ANOVA with Games– Howell correction. Non-normally distributed data were analysed using the Wilcoxon rank-sum test for two groups or Kruskal–Wallis test with Dunn’s post hoc for multiple groups.

### NF-kB nuclear translocation assay

HPCs (1.2 × 10⁴ cells per well) were seeded in laminin-coated 96-well plates. The following day cells were treated ± 10 ng/mL TNF-α and fixed after 0, 15, 30, 45, and 60 min, and 2, 3, 4, 24, and 48 h. Cells were immunostained for NF-κB p65 and counterstained with DAPI. High-content imaging was performed on an Opera Phenix (PerkinElmer), and images were analysed using Harmony software (PerkinElmer). Nuclear and cytoplasmic NF-κB p65 signal intensities were measured, and the nuclear fraction of p65 fluorescence was calculated as described below:

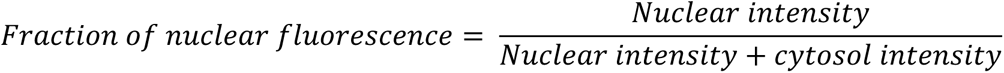

### Cytokine and chemokine secretion analysis

HPCs (1.2 × 10⁵ cells) were seeded in laminin-coated 24-well plates (Thermo Fisher Scientific, Cat. No. 142475). The following day, the media was replaced with proliferation medium ± TNF-α. After 24 h and 48 h, supernatants were collected and stored at –80 °C. For differentiation, the cells cultured in proliferation media for 48 hours were washed twice with differentiation medium for 30 min, then cultured in differentiation medium ± TNF-α. Supernatants were collected after 24 h, 48 h, and 7 d. IL-6 levels were quantified using a human IL-6 ELISA Kit (BioLegend, Cat. No. 430504). A 13-plex panel of human pro-inflammatory chemokines (Supplementary Table 6) was analysed using a multiplex bead-based assay kit (BioLegend, Cat. No. 740985). Data was acquired using a FACSCanto II (BD Biosciences) flow cytometer and analysed using the LEGENDplex data analysis software suite (BioLegend).

### Differentiation assay for immunocytochemistry

HPCs were seeded at 1.2 × 10⁴ cells per well in laminin-coated 96-well plates (Thermo Fisher Scientific, Cat. No. 140675) in proliferation medium containing 4-OHT, bFGF, and EGF. The following day, the medium was replaced with proliferation medium ± TNF-α. After 48 h, cells were washed twice with differentiation medium (no 4-OHT, bFGF, or EGF) for 30 min, then cultured in differentiation medium ± TNF-α for 7 d, with media changes every 48 h. In a subsequent experiment, cells were treated with ± 1 ng/mL TNF-α and ± 10 µg/mL anifrolumab.

### Neurite outgrowth analysis

Neurite outgrowth of DCX+ and MAP2+ cell populations was assessed using Harmony (PerkinElmer). Cells were identified as described in the High-Content Imaging and Analysis section. Neurites were detected using the ‘Find Neurites’ building block, and neurite morphology parameters, including the length of the longest neurite, were quantified.

### Cell culture and treatment for the single-cell RNA sequencing

#### Proliferation

HPCs were seeded into 20 μg/ml mouse laminin (Sigma, Cat. No. L2020) coated T175 flasks (Sigma, Cat. No. L2020) at 2.9 × 10⁶ cells for control or low TNF-α (0.1 ng/mL) conditions, and 5 × 10⁶ cells for high TNF-α (1 ng/mL) conditions. The following day, the media was replaced with proliferation medium ± TNF-α. The cells were harvested after 48 h.

#### Differentiation

HPCs were cultured as described above in proliferation medium ± TNF-α for 48 hrs. Next, the media was changed to differentiation media ± 0.1 ng/ml TNF-⍺ or 1 ng/ml TNF-⍺. The HPCs were allowed to differentiate for seven or fourteen days. To mimic a chronic TNF-⍺-mediated proinflammatory environment the differentiation media was changed every 48 hours to fresh differentiation media ± 0.1 ng/ml TNF-⍺ or 1 ng/ml TNF-⍺. After seven or fourteen days of differentiation the cells were harvested.

### Harvesting cells for single-cell RNA sequencing

On the day of harvest, medium was removed and cells washed with pre-warmed DPBS (Gibco, Cat. No. 14190144). Cells were detached using pre-warmed Accutase (Gibco, Cat. No. A1110501) at 37 °C, 5% CO₂ for 5–10 min, harvested with pre-warmed DMEM/F12 (Gibco, Cat. No. 21331020), and centrifuged at 100 × g for 5 min. Pellets were gently resuspended in 1 mL of the warm proliferation or differentiation medium. Viability was confirmed at ≥ 85% using Trypan Blue (Thermo Fisher Scientific, Cat. No. 15250061,) and a hemocytometer before proceeding to antibody labelling.

### Cell surface staining with hashtag oligonucleotide antibodies

For 3′ scRNA-seq multiplexing on the 10x Genomics Chromium system, samples were labelled with TotalSeq™-B anti-human hashtag oligonucleotide (HTO) antibodies targeting CD298 and β2-microglobulin (BioLegend: Hashtag 1 Cat. No. 394631; Hashtag 2 Cat. No. 394633; Hashtag 3 Cat. No. 394635). Optimal antibody concentration (1:5,000; 125 ng/mL) was determined by titrating PE-conjugated anti-CD298 (BioLegend, Cat. No. 341704) and anti-B2M (BioLegend, Cat. No. 316305) at a 1:1 equimolar ratio, followed by ICC to confirm 100% labelling with minimal background. Samples were individually labelled with HTOs prior to pooling at the sequencing facility.

1 × 10⁶ cells per sample were pelleted (300 × g, 5 min, 4 °C) and resuspended in 45 µL Cell Staining Buffer (CSB) (BioLegend, Cat. No. 420201). Cells were blocked with 5 µL TruStain FcX™ (BioLegend, Cat. No. 422301) for 10 min at 4 °C. TotalSeq™-B HTO antibodies were diluted 1:2,500 in CSB and mixed 1:1 with blocked cells to reach a final concentration of 1:5,000. Samples were incubated for 30 min at 4 °C. Cells were washed 3× in CSB (300 × g, 5 min, 4 °C) resuspending using wide-bore pipette tips (Starlab, Cat. No. E1011-9618). Final concentration was adjusted to 700–1,200 cells/µL, assuming ∼50% cell loss. Cell suspensions were filtered through 40 µm Flowmi™ Cell Strainers (Bel-Art, Cat. No. H13680-0040), and viability (>80%) was confirmed before proceeding to the 10x Genomics workflow. All steps were performed on ice.

### Single-cell capture, cDNA library preparation, and sequencing

HTO-labelled samples were pooled and processed by the Genomics Research Platform at Guy’s & St Thomas’s Hospital (London, UK). The differentiation samples (three pools) and the proliferation samples (one pool) were delivered to the facility on separate days. Equal cell numbers were combined, and 18,000 cells per pool were loaded onto separate lanes of the 10x Chromium X instrument (10x Genomics) for droplet-based single-cell capture. cDNA libraries were prepared using the Chromium Single Cell 3′ Reagent Kit v3 (10x Genomics, Cat. No. PN-1000268,). Libraries were sequenced on an Illumina NextSeq 2000 system using 50 bp paired-end reads.

### Read alignment and sample demultiplexing

FASTQ files from gene expression and HTO libraries were processed using Cell Ranger v7.1 (10x Genomics) with the cellranger multi pipeline. Data were imported into R (v4.3.2) using Seurat v5 via the Read10X() function, returning UMI and HTO count matrices. Cells with barcodes present in both matrices were retained, and RNA counts were used to create a Seurat object (CreateSeuratObject()), with HTO data added as an independent assay (CreateAssayObject()). HTO data were normalised using centred log-ratio transformation and demultiplexed using HTODemux(assay = ‘HTO’) with default parameters. Only singlets with confident HTO classification were retained for downstream analysis.

### Quality control

QC was performed separately on merged Seurat objects for differentiation and proliferation samples. At the cell level, cells were excluded if they had >15% mitochondrial gene expression, a novelty score (log₁₀[genes/UMIs]) <0.8, <800 or >8,000 genes, or <1,000 or >45,000 transcripts. This yielded 22,133 high-quality cells (15,053 derived from differentiation samples; 7,080 derived from proliferation samples). At the gene level, genes not expressed in any cells or expressed in ≤5 cells were excluded. Post-filtering, 26,288 genes were retained in the differentiation dataset and 21,951 in the proliferation dataset.

### Normalisation, dimensionality reduction, and clustering

The differentiation and proliferation Seurat objects were merged and normalised using sctransform (v2) from Seurat. Dimensionality reduction was performed using principal component analysis (PCA) on the top 3,000 highly variable genes. An ElbowPlot indicated that the first 50 principal components captured the majority of variance in the dataset which was used for downstream analysis. We constructed a shared nearest neighbour graph using FindNeighbors(), tested clustering across resolutions from 0.1 to 1.2 (FindClusters()), and visualised cellular heterogeneity via UMAP (RunUMAP(dims = 1:50)), displayed with DimPlot(reduction = “umap”).

### Cluster annotation

Clusters were initially over-resolved using high-resolution settings and subsequently merged if they lacked distinct marker genes, were not biologically meaningful, or shared multiple canonical markers. This iterative approach—clustering, merging, and re-clustering—was guided by marker gene expression. Cluster markers were identified using the FindAllMarkers() function in Seurat, applying Wilcoxon rank-sum testing to compare each cluster against all others. Marker genes were required to be expressed in ≥25% of cells within the cluster and show positive differential expression (log₂ fold change ≥0.25). Ultimately, 15 clusters were identified and annotated based on culture condition, treatment, cell cycle stage, and canonical marker expression.

### Functional enrichment analysis

^111^Gene Ontology (GO) over-representation analysis was performed using the ClusterProfiler package^83^. The top 100 differentially expressed genes (ranked by log fold change) between clusters were used as input. Raw p-values < 0.05 were considered for enrichment. Multiple testing correction was performed using the Benjamini–Hochberg method, with the default adjusted p-value threshold of 0.2.

### Transcription factor activity inference

Transcription factor (TF) activity was inferred using decoupleR (v2.8)^84^. CollecTRI^85^ was retrieved via OmnipathR^86^. The univariate linear model generates a TF enrichment score by fitting a linear model of gene expression based on TF–gene interaction weights. The t-value of the slope (ULM score) indicates TF activity: positive scores denote activation and negative scores indicate inactivity.

### Gene module scoring for NSC-specific type I IFN response signature

Cells were scored using a 300-gene NSC-specific type I IFN response signature derived from murine SVZ NSCs treated *ex vivo* with IFN-β^38^. Average gene set expression per cell was calculated using the Seurat function AddModuleScore().

### Gene module scoring for human immature dentate granule cell signature

Cells were scored against the top 20 genes from a gene list identifying human immature dentate granule cells^9^. Average gene set expression per cell was calculated using the AddModuleScore() funtion.

### Conditioned media experiment for immunocytochemistry

To generate conditioned media (CM) 3 x 10^5^ HPCs were seeded into six well plates coated with 20 μg/ml mouse laminin (Sigma, Cat. No. L2020). The following day, the media was replaced with fresh media ± 0.1 ng/ml, 1 ng/ml, or 10 ng/ml TNF-⍺. After 24 hours, media was switched to control media without TNF-⍺ and cells were cultured for an additional 24 hours before CM collection. This approach captured HPC-secreted factors in response to TNF-⍺ while eliminating residual TNF-⍺. To assess CM effects on IFN-regulated protein ISG15 expression, HPCs were seeded into laminin-coated 96-well plates. The next day, media was replaced with 70% CM mixed with 30% fresh media to maintain cell viability. After 48 hours, cells were fixed, immunostained, and imaged to quantify the percentage of ISG15⁺ HPCs.

### Protein synthesis inhibition

To investigate the requirement of *de novo* protein synthesis in the TNF-⍺-induced upregulation of candidate genes, we utilised the global protein translation inhibitor, cycloheximide. The HPCs were treated as described in the results section with 10 μg/ml cycloheximide (Merck, Cat. No. 01810).

### Blocking NF-kB signalling

Cells were seeded in six-well plates coated with 20 μg/ml mouse laminin (Sigma, Cat. No. L2020). After 24 hours, the media was replaced with media containing 10 μM JSH23 (Abcam, Cat. No. ab144824) dissolved in 100% dimethyl sulfoxide (DMSO) or with DMSO as control. The final DMSO concentration in both groups was 0.1%. Following a 30-minute pre-treatment, 1 ng/ml TNF-⍺ was added as indicated in the results section, and cells were harvested for gene expression analysis.

### Probing the dependency of IFNAR in TNF-⍺-induced type I IFN signalling

To assess IFNAR dependency in TNF-⍺-induced type I IFN signalling, HPCs (1.2 × 10^4) were seeded in laminin-coated 96-well plates. The following day, cells were treated with or without 10 μg/ml anifrolumab (Bio X Cell, Cat. No. SIM0022), an anti-IFNAR1 antibody, or isotype control (Bio X Cell, Cat. No. SIM0014), and with or without 1 ng/ml TNF-⍺. After 24 h, cells were fixed and stained for ISG15, STAT1, and DAPI. High-content imaging quantified ISG15+ and STAT1+ cells as described.

### Blocking JAK activity

1.2 × 10^4^ HPCs were seeded in laminin-coated 96-well plates (Thermo Fisher Scientific, Cat. No. 167008). The following day, cells were treated with or without 16 nM JAK inhibitor I (Cayman Chemical, Cat. No. CAY15146) and with or without 1 ng/ml TNF-⍺. After 24 h, media was removed, cells fixed and stained, and high-content imaging was used to quantify ISG15+ HPCs.

### Evaluating cell surface protein expression with flow cytometry

Cells were cultured in laminin-coated T75 flasks (Sigma, Cat. No. L2020) and treated ± 1 ng/ml TNF-⍺ for 24 h. Cells were detached with Accutase (Sigma, Cat. No. A1110501) for five min in the incubator, harvested, centrifuged, and resuspended in PBS. Cells were stained with Zombie NIR viability dye (BioLegend, Cat. No. 423105; 1:2000 in PBS) for 15 min at room temperature. Cells were washed and stained with antibodies (Supplementary Table 7) at 1:100 in staining buffer (PBS + 10 mM HEPES, 2 mM EDTA, 0.5% FCS) for 40 min at 4 °C. After washes, cells were resuspended in staining buffer. Data were acquired on a BD FACS Canto II using FACSDiva v9.2 and analysed with FlowJo v10.10.

### Isolating peripheral blood mononuclear cells from leukocyte cones

Peripheral blood mononuclear cells (PBMCs) were isolated from leukocyte cones obtained from three healthy donors via NHS Blood and Transplant (NHSBT, Tooting, UK). Donors provided generic consent for research use. Leukocyte cones were cleaned with ethanol and processed in a biosafety cabinet. Blood was drained into a 50 ml tube which was topped up to 50 ml with PBS containing 2% FCS. After gently mixing it was centrifuged at 800 × g for 10 minutes. The pellet was resuspended 1:1 in DPBS and layered onto 15 ml Ficoll-Paque PLUS (Cytiva, Cat. No. 17144002). After centrifugation at 450 × g for 30 minutes with minimal acceleration and no brake, the buffy coat was collected, washed twice with cold DPBS, and centrifuged at 400 × g for 10 minutes at 4°C. A final centrifugation step at 200 × g for 10 minutes at 4°C was performed to reduce platelet contamination. Cells were counted, resuspended in freezing media (90% FBS, 10% DMSO) at 360–600 × 10^6^ cells/ml, aliquoted into cryovials, frozen overnight at −80°C in a Mr. Frosty container, and transferred to liquid nitrogen for long-term storage. Each vial contained 9–15 × 10^6^ cells.

### Chemotaxis assay

#### T cell activation and expansion protocol

PBMC cryovials were thawed and seeded in T75 flasks at 37°C with 5% CO₂ in complete RPMI (10% FCS (Thermo Fisher Scientific, Cat. No. 26140079), 1% penicillin-streptomycin (Thermo Fisher Scientific, Cat. No. 15140122)). After 24 hours, nonadherent lymphocytes were collected, centrifuged, and cultured in complete RPMI supplemented with 100 IU/ml recombinant human IL-2 (BioLegend, Cat. No. 689104) and 2.5 μg/ml phytohemagglutinin-L (PHA-L) (Thermo Fisher Scientific, Cat. No. 00-4977-93) in fresh T75 flasks. After 48 hours, cells were harvested, washed, and resuspended in complete RPMI with 100 IU/ml IL-2. The cells were cultured for a total of 10 days and were resuspended in fresh complete RPMI media supplemented with 100 IU/ml IL-2 every other day.

#### Transwell chemotaxis assay

Transwell migration assays were performed using 96-well transwell plates with 5 μm pore membranes (Corning, Cat. No. 3387). Upper wells were coated overnight with 2% BSA in PBS to prevent cell adhesion and washed thrice with PBS the following day. Activated lymphocytes were harvested, washed, and resuspended at 4–5 × 10^6 cells/ml in migration media (complete RPMI diluted 1:10 in neat RPMI), HPC proliferation media, or HPC differentiation media as appropriate. For CXCR3 dependency assays, cells were pretreated with 1 μM AMG487 (Cayman Chemical, Cat. No. 28416) or DMSO control for 30 minutes at 37°C with rotation. 75 μl cell suspension were added to upper chambers. Lower chambers were filled with 235 μl media containing 100 ng/ml recombinant human CXCL10 (R&D Systems, Cat. No. 266-IP), TNF-⍺-conditioned media (CM) from proliferating or differentiating HPCs, or CM from HPCs treated ± TNF-⍺, JAK inhibitor I, or anifrolumab. After 3 hours at 37°C, migrated cells in the lower chamber were collected into 96-well plates for analysis.

#### Flow cytometry of migrated cells

Migrated cells were collected by spinning the plate at 2000 rpm for 1 min and washed with 200 μl DPBS. Cells were stained with Zombie NIR viability dye (BioLegend, Cat. No. 423105; 1:2000 in DPBS) for 15 min at room temperature in the dark, followed by quenching with 100 μl DPBS and centrifugation. For CXCR3 surface expression, cells were incubated with anti-CXCR3 antibody (1:100 in staining buffer: DPBS + 10 mM HEPES + 2 mM EDTA + 0.5% FCS) at 37°C for 15 min. Next cells were stained with a master mix of antibodies (Supplementary Table 8; 1:50) for 40 min at 4°C. Cells were washed twice with staining buffer and resuspended in 100 μl staining buffer. Data were acquired on a BD FACS Canto II with high-throughput sampler using FACSDiva v9.2 (BD Biosciences) with settings: flow rate 3.0, sample volume 50 μl, mixing volume 50 μl, mixing speed 200, two washes with 200 μl wash volume. Analysis was performed using FlowJo v10.10.

#### Generating conditioned media from HPCs for chemotaxis study

HPCs (3.5 × 10⁵) were seeded into six-well plates coated with 20 μg/ml mouse laminin (Sigma, Cat. No. L2020). After 24 h, media was replaced with proliferation media ± 1 ng/ml TNF-⍺. After 48 h, CM was collected and centrifuged at 1500 × g for 10 min at 4°C. Supernatants were transferred to new tubes to remove cell debris and stored at –80°C until use.

#### Generating conditioned media from differentiating HPCs

After 48 h of treatment in proliferation conditions as described above, media was replaced with differentiation media ± 1 ng/ml TNF-⍺. Cells were differentiated for 7 days, with media ± 1 ng/ml TNF-⍺ refreshed every 48 h to simulate chronic inflammation. At the end of the differentiation period, CM was collected, centrifuged at 1500 × g for 10 min at 4°C, and the supernatants were transferred to new tubes and stored at –80°C for chemotaxis assays.

#### Generating conditioned media from HPCs blocking JAK activity or IFNAR

3.5 × 10⁵ HPCs were seeded in 6-well plates coated with 20 μg/ml mouse laminin (Sigma, Cat. No. L2020). The following day, media was replaced with fresh proliferation media with either DMSO (vehicle control), 1 ng/ml TNF-⍺, 10 μg/ml anifrolumab (anti-IFNAR1; Bio X Cell, Cat. No. SIM0022) ± 1 ng/ml TNF-⍺, or 0.15 nM JAK inhibitor I (Cayman Chemical, Cat. No. CAY15146) ± 1 ng/ml TNF-⍺ After 24 h, CM was collected, centrifuged at 1500 × g for 10 min at 4°C, and supernatants were transferred to fresh tubes and stored at –80°C for chemotaxis assays.

## Data availability

All source data will be made available via the repository platform *figshare.com* and scripts on *github.com* at the same time as the peer-reviewed publication.

## Authors contribution

L.S.K. and S.T. contributed equally and directed the research. T.A.D.N., L.S.K., and S.T. designed the experiments. T.A.D.N. wrote the manuscript with contributions from L.S.K. T.A.D.N. performed the majority of the experiments and conducted data analysis. T.A.D.N. and D.T.R. generated the scRNA sequencing data in collaboration. A.B. collected and analysed the RT-qPCR data. S.S. performed the morphological analysis of differentiated HPCs. H.L. helped design and generate the differentiation and chemokine data. T.A.D.N. and L.A.O’N. generated the ELISA and chemokine data. S.J. advised on STAT signalling and helped generate the STAT western blot data. All authors read and revised the manuscript.

## Acknowledgements

T.A.D.N. was funded by the Wellcome Trust as part of the “Neuro-Immune Interactions in Health & Disease Wellcome Trust PhD Programme [218452/Z/19/Z]”. The work was supported by a grant awarded to S.T. by the Medical Research Council UK [MR/S00484X/1] and awards to L.S.K. by the Medical Research Council Discovery Award [MR/N012345/1]. D.T.R. was supported by a PhD Studentship awarded by the Medical Research Council UK [MR/R015643/1], H.L. was supported by a grant from The Galen and Hilary Weston Foundation awarded to S.T. Some panels of Figure 5 and 6 were created with Biorender.com with a licence to S.T. at King’s College London.

We thank George Chennell and Chen Liang (Wohl Cellular Imaging Centre, King’s College London) for technical support with high-content imaging; the Genomics Research Platform (Guy’s and St Thomas’ NHS Foundation Trust) for sequencing support; Franziska Denk for advice on scRNA sequencing design; Robert Köchl and Aritz Lasarte Cía for input on chemotaxis assays; Josephine Eum for guidance on PBMC isolation; and Daniel Davies for advice on flow cytometry staining.

## Ethics declaration

The authors declare that ethical standards prevailing at their institution have been observed throughout this study.

## Extended data

**Extended Fig. 1.**
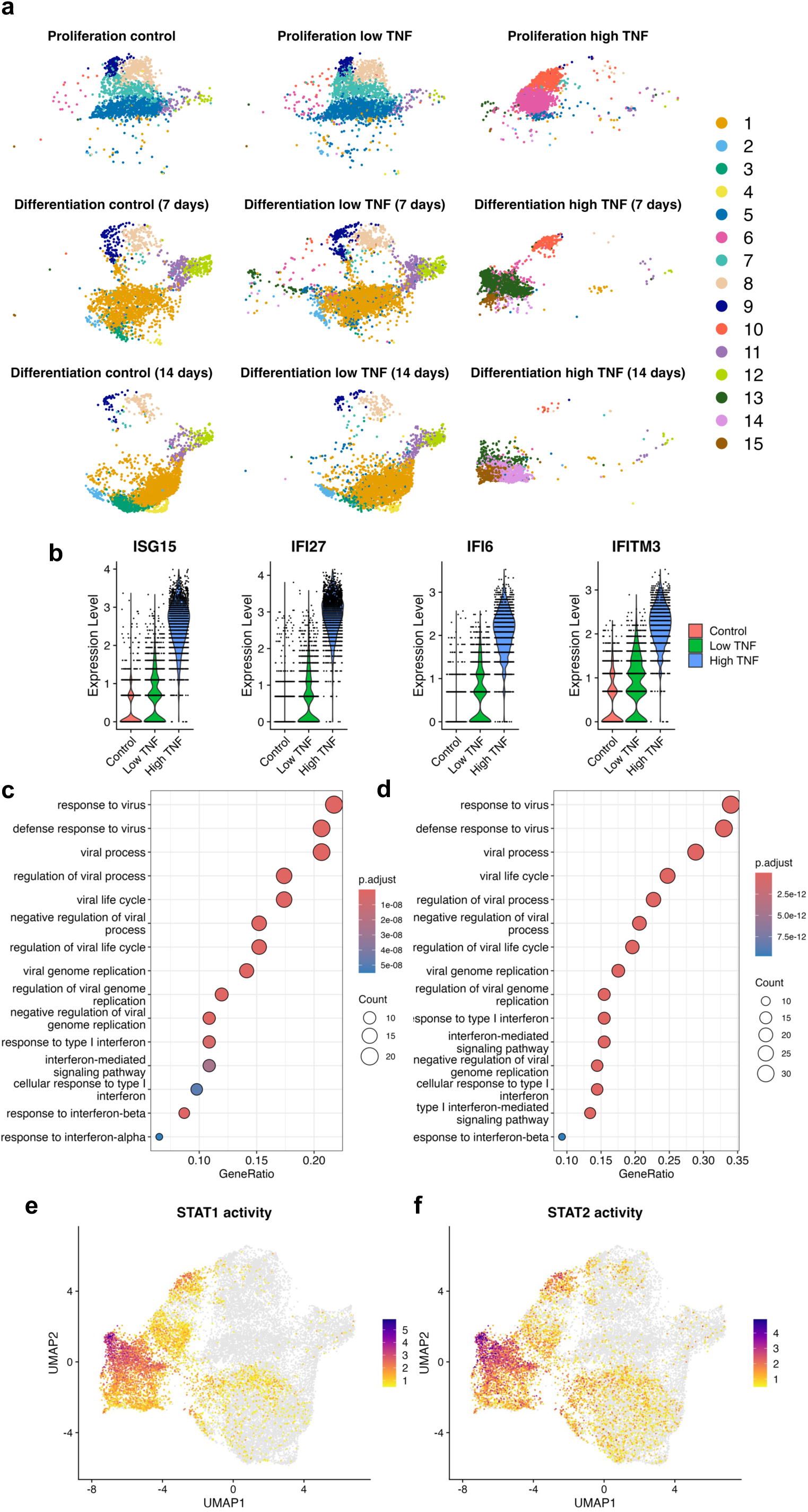
TNF-⍺ drives the upregulation of type I IFN genes in HPCs. **a**, UMAP plot showing transcriptional clustering of single cells split by samples. Colours indicate the 15 cell clusters identified. **b**, Violin plots showing dose-dependent expression of the type I IFN stimulated genes ISG15, IFI27, IFI6, and IFITM3 in the HPCs treated for 48 hrs with vehicle (red), 0.1 ng/ml (low, green), or 1 ng/ml (high, blue) TNF-⍺. **c**, Dot plot showing gene ontology (GO) enrichment analysis of biological processes terms based on the top 100 genes upregulated genes in cluster 5 as compared to cluster 6. **d**, Dot plot showing gene ontology (GO) enrichment analysis of biological processes terms based on the top 100 genes upregulated genes in cluster 10 as compared to cluster 7, 8, and 9. **e**, UMAP plot showing the predicted transcription factor activity of STAT1. **f**, UMAP plot showing the predicted transcription factor activity of STAT2.

**Extended Fig. 2.**
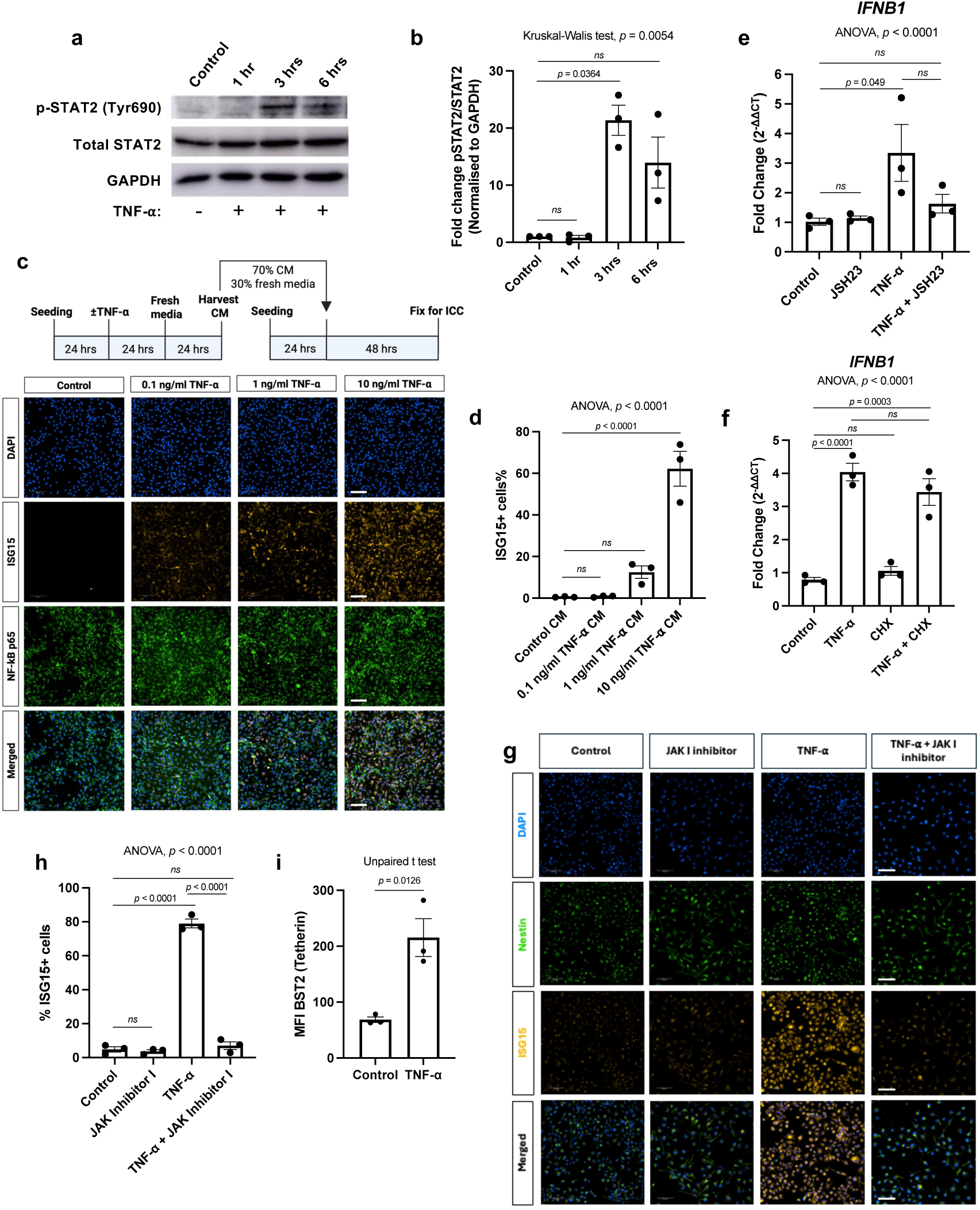
TNF-⍺ drives type I IFN signalling in human HPCs via autocrine/paracrine signalling via IFNAR. **a**, Representative western blots of three independent experiments showing the activation of STAT2 in HPCs treated with 1 ng/ml TNF-⍺ for 1 h, 3 h, 6 h or 12 h. Cell lysates were probed for phosphorylated STAT2 (p-STAT2, Tyr690), total STAT2, and GAPDH as a loading control. **b**, Quantification of STAT2 activation in human hippocampal progenitor cells from A. pSTAT2 and STAT2 intensities were normalised to GAPDH intensity before calculating the fold change pSTAT2/STAT2 relative to control. Data represent mean ± SEM from three independent experiments. Statistical analysis: Kruskal-Wallis test followed by Dunn’s post hoc correction. **c**, Representative images of three independent experiments with similar results showing the expression of ISG15 (orange) on HPCs. Scale bar = 100 µm. HPCs treated for 48 hrs with conditioned media (CM) from other HPCs treated ± TNF-⍺ for 24 hrs followed by 24 hrs treatment with media. **d**, Quantification of the percentage of ISG15+ cells based on C. Data represent mean ± SEM. Statistical analysis: one-way ANOVA followed by Bonferroni’s multiple comparisons test. **e**, RT-qPCR data showing the relative gene expression of *IFNB1* in HPCs pre-treated with 10 µM JSH23 or DMSO, followed by three hours of treatment ± 1 ng/ml TNF-⍺. Data represent as mean ± SEM of three independent experiments. Statistical analysis: one-way ANOVA followed by Bonferroni’s multiple comparisons test. **f**, RT-qPCR data showing the relative gene expression of *IFNB1* in HPCs pre-treated with 10 µg/ml cycloheximide (CHX) or DMSO followed by three hours of treatment ± 1 ng/ml TNF-⍺. Data represent as mean ± SEM of three independent experiments. Statistical analysis: one-way ANOVA followed by Bonferroni’s multiple comparisons test. **g**, Representative images of three independent experiments with similar results showing the expression of ISG15 (orange) on HPCs. Scale bar = 100 µm. HPCs were treated for 24 hrs with 16 nM JAK inhibitor I or DMSO, ± 1 ng/ml TNF-⍺. **h**, Quantification of the percentage of ISG15+ cells based on G. Data represent mean ± SEM. Statistical analysis: one-way ANOVA followed by Bonferroni’s multiple comparisons test. **i**, Cell surface expression of IFN-regulated BST2 (tetherin) quantified by measuring the mean fluorescence intensity (MFI) on HPCs treated 24 hrs ± 1 ng/ml TNF-⍺ using flow cytometry. Data represent mean ± SEM. Statistical analysis: unpaired t-test.

## Supplementary Information

**Supplementary Fig. 1.**
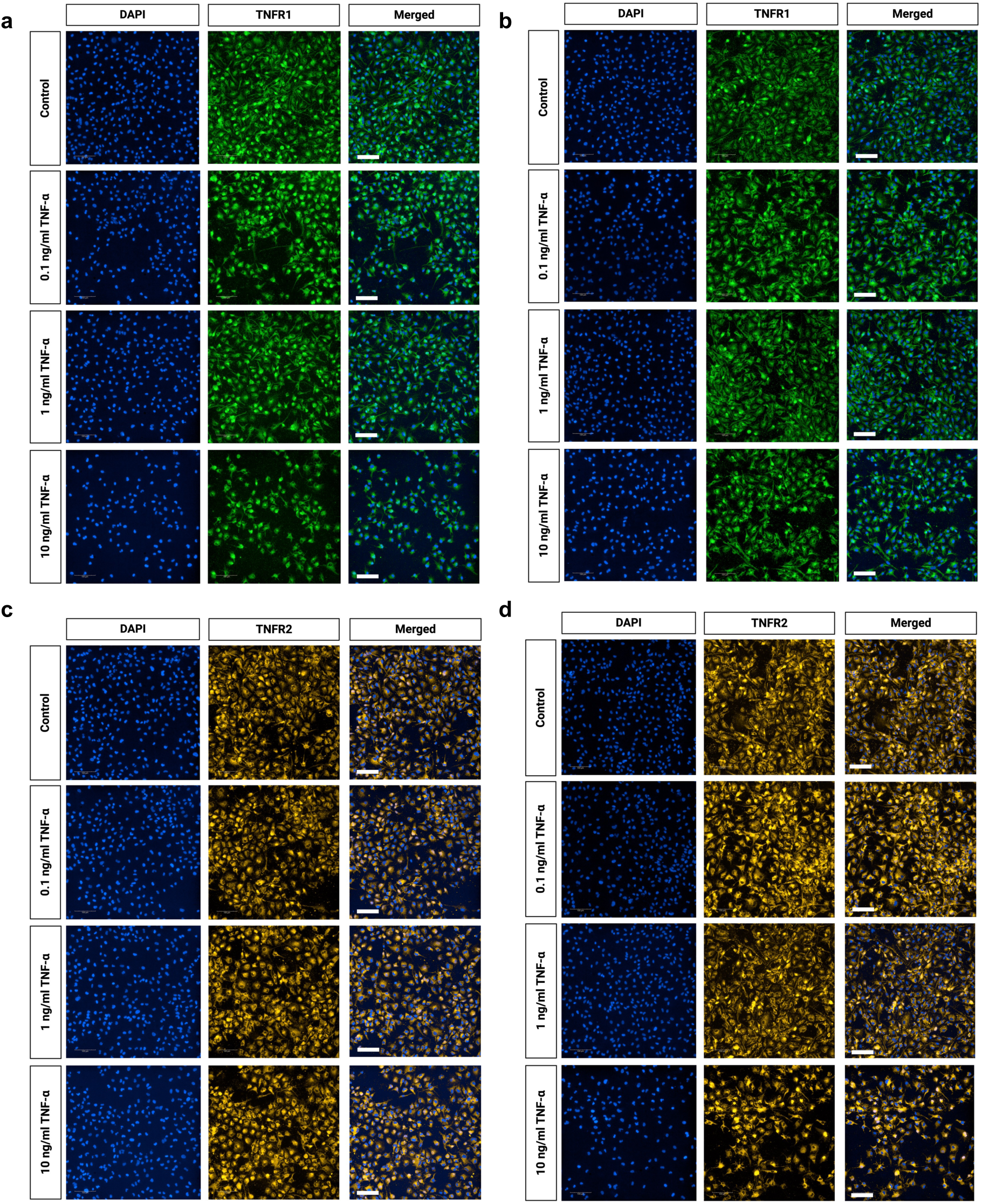
TNFR expression on human HPCs treated ± TNF-⍺. **a,** and **b** Representative images of three independent experiments with similar results showing the expression of TNFR1 (green) on HPCs treated ± TNF-⍺ for **a** 24 h or **b** 48 h. **c** and **d** Representative images of three independent experiments with similar results showing the expression of TNFR2 (orange) on HPCs treated ± TNF-⍺ for **c** 24 h or **d** 48 h. Scale bar = 100 μm. DAPI; 4′,6-diamidino-2-phenylindole, HPC; hippocampal progenitor cells, TNFR1; tumour necrosis factor receptor 1.

**Supplementary Fig. 2.**
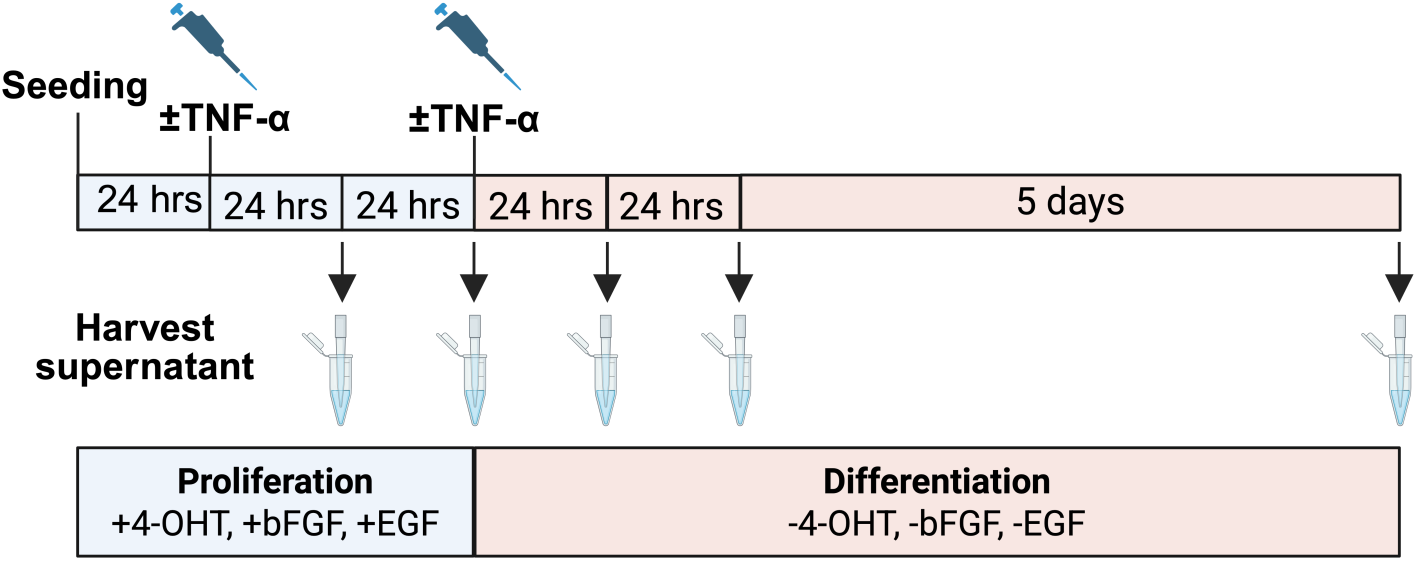
Schematic overview of the experimental design for harvesting supernatant from HPCs and their differentiated progeny. The day following seeding, the proliferating HPCs were treated ± TNF-⍺ (0.1 ng/ml, 1 ng/ml, or 10 ng/ml). After 24 h, the supernatant was harvested. After 48 h of treatment, the supernatant was harvested, and the media was changed to induce differentiation (removal of 4-OHT, bFGF, and EGF) ± TNF-⍺ (0.1 ng/ml, 1 ng/ml, or 10 ng/ml). The supernatant of the differentiating cells was harvested after 24 h, 48 h, or 7 days of differentiation. 4-OHT; 4-hydroxytamoxifen, bFGF; basic fibroblast growth factor, EGF; epidermal growth factor.

**Supplementary Fig. 3.**
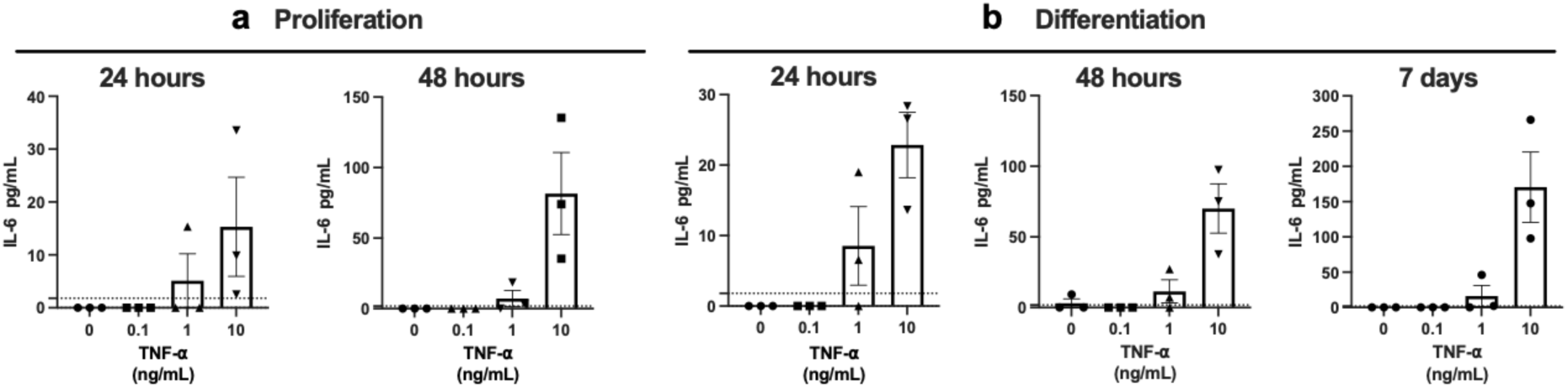
ELISA measurement of IL-6 in the supernatant of HPCs and their differentiated progeny treated with TNF-⍺. **a,** Concentrations of IL-6 in the supernatant of proliferating HPCs 24 h and 48 h after TNF-⍺ treatment. **b,** IL-6 in the supernatant of differentiating HPCs 24 h, 48 h, and 7 days after TNF-⍺ treatment. Data is presented as mean ± SEM of three independent experiments.

**Supplementary Fig. 4.**
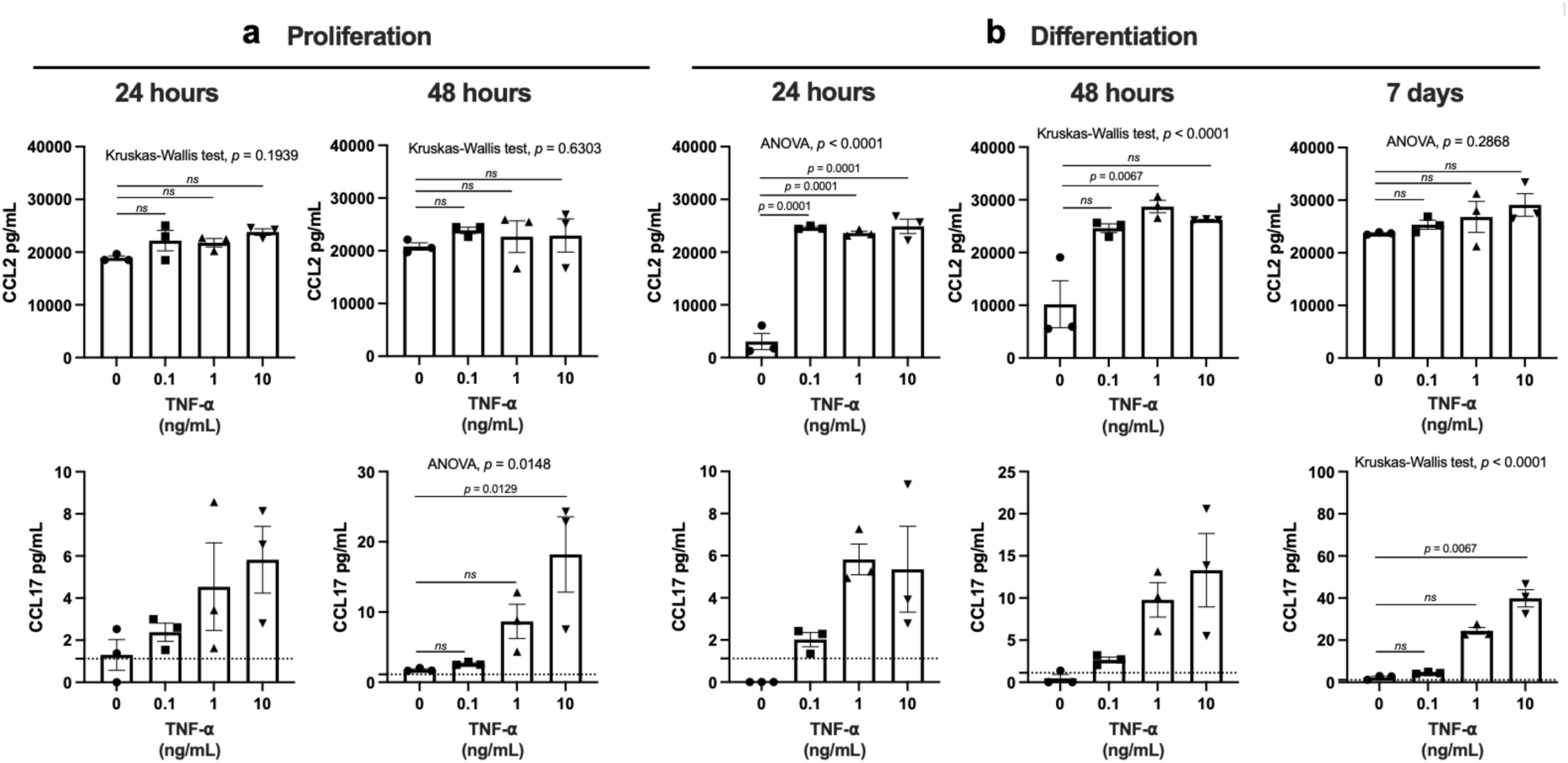
CCL2 and CCL17 levels in the supernatant of HPCs and their differentiated progeny treated with TNF-⍺. **a,** Concentrations of CCL2 (upper panel) and CCL17 (lower panel) in the supernatant of proliferating HPCs 24 h and 48 h after TNF-⍺ treatment. **b,** CCL2 (upper panel) and CCL17 (lower panel) in the supernatant of differentiating HPCs 24 h, 48 h, and 7 days after TNF-⍺ treatment. Data is presented as mean ± SEM of three independent experiments. Statistical test: Kruskal-Wallis test followed by Dunn’s post hoc correction, or one-way ANOVA followed by Bonferroni’s multiple comparisons, as indicated above the graphs. Dashed lines mark the detection limit for CCL17 (1.12 pg/ml).

**Supplementary Fig. 5.**
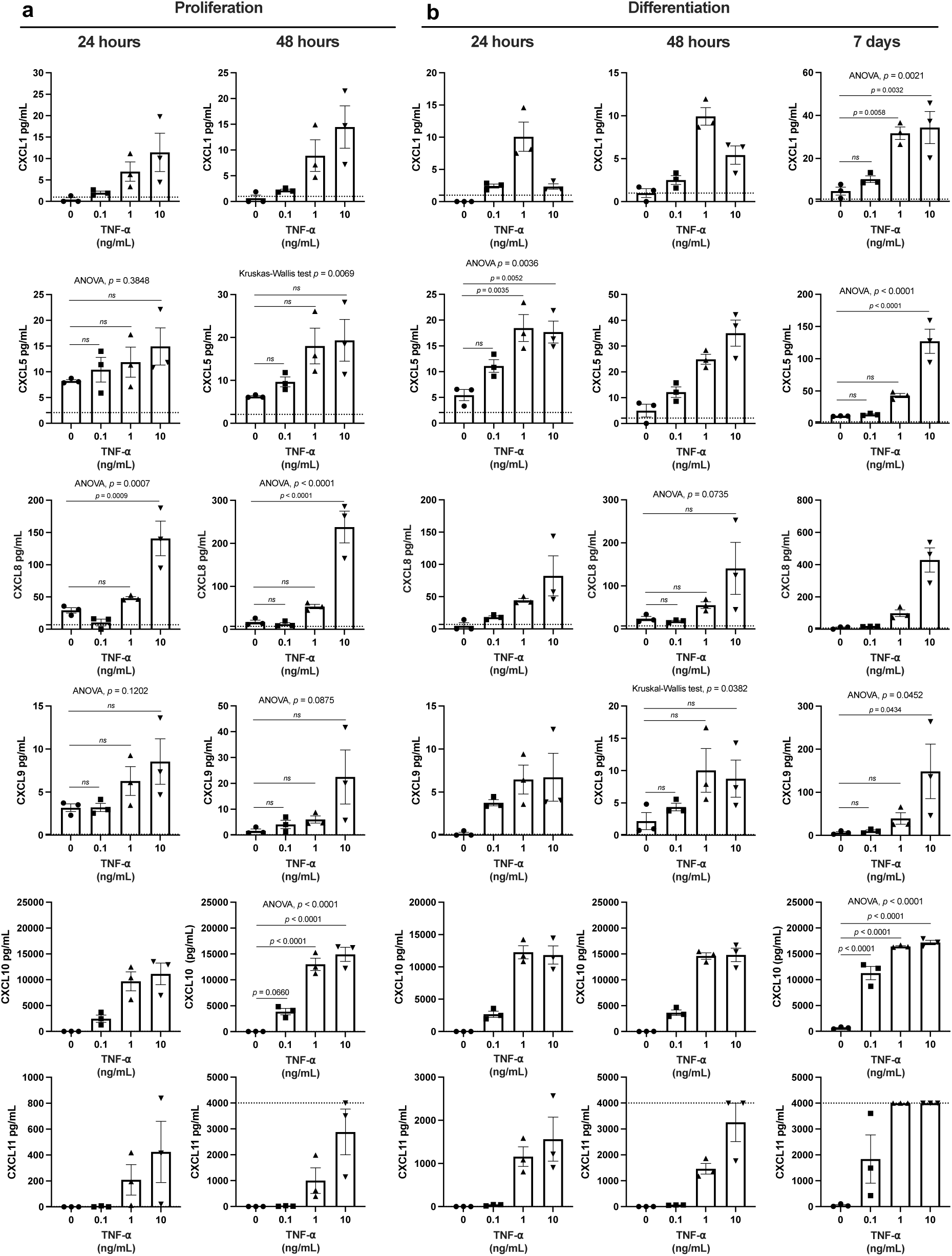
CXC chemokine levels in the supernatant of HPCs and their differentiated progeny treated with TNF-⍺. **a,** Concentrations of CXCL1, CXCL5, IL-8, CXCL9, CXCL10, and CXCL11 in the supernatant of proliferating HPCs 24 h and 48 h after TNF-⍺ treatment. **b,** CXCL1, CXCL5, IL-8, CXCL9, CXCL10, and CXCL11 levels in the supernatant of differentiating HPCs 24 h, 48 h, and 7 days after TNF-⍺ treatment. Data is presented as mean ± SEM of three independent experiments. Statistical test: Kruskal-Wallis test followed by Dunn’s post hoc correction or one-way ANOVA followed by Bonferroni’s multiple comparisons as indicated above the graphs. Dashed lines mark the detection limits (lower limit for CXCL1: <0.99 pg/ml, lower limit for CXCL5: <2.07 pg/ml, lower limit for CXCL8: <6.86 pg/ml, lower limit for CXCL9: <0.098 pg/ml, lower limit for CXCL10: <23.44 pg/ml, lower limit for CXCL11 <2.13 pg/ml, and upper limit for CXCL11 > 4000 pg/ml). For visualisation, data points below the detection limit were set to zero, and data points above the upper detection limit for CXCL11 were set to the limit of 4000 pg/ml.

**Supplementary Fig. 6.**
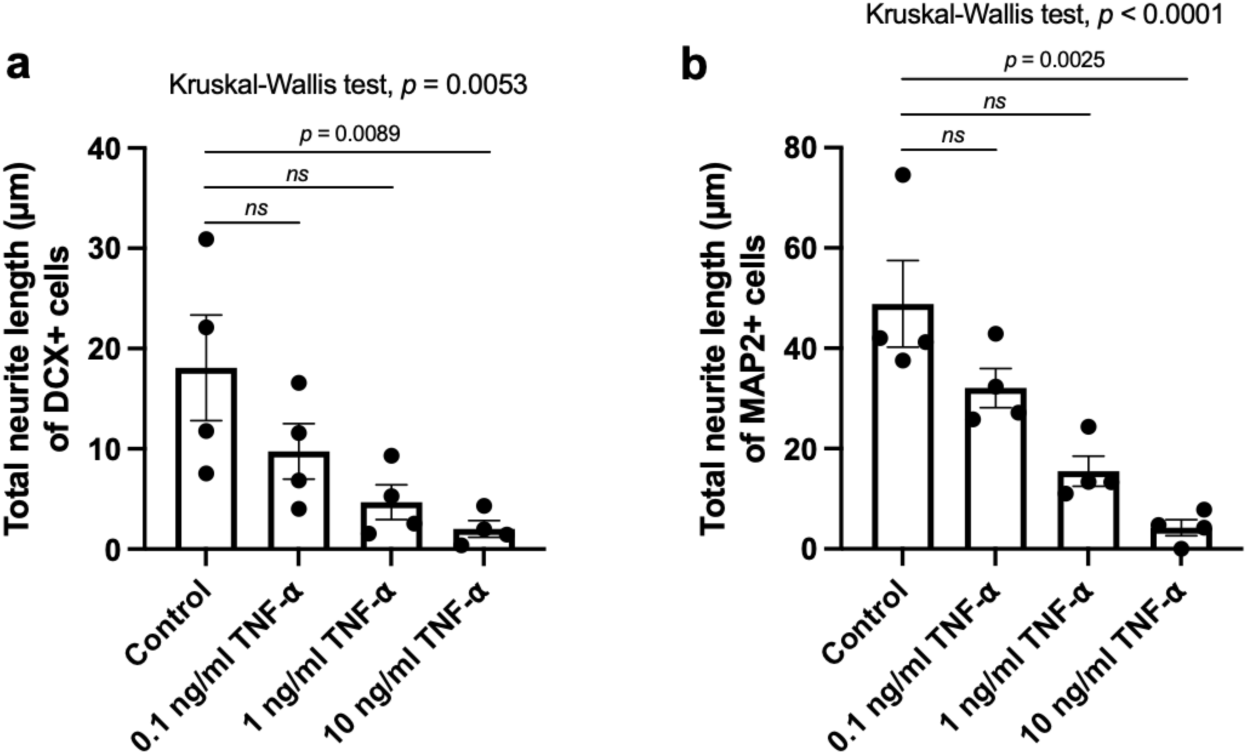
Chronic TNF-⍺ dose-dependently decreases the neurite length of DCX+ and MAP2+ differentiating HPCs. Quantification of the total neurite length (µm) of **a** DCX+ cells and **b** MAP2+ cells after seven days of differentiation ± chronic treatment with 0.1, 1, or 10 ng/ml TNF-⍺. Data represent mean ± SEM from four independent experiments. Statistical analysis: Kruskal-Wallis test followed by Dunn’s post hoc correction.

**Supplementary Fig. 7.**
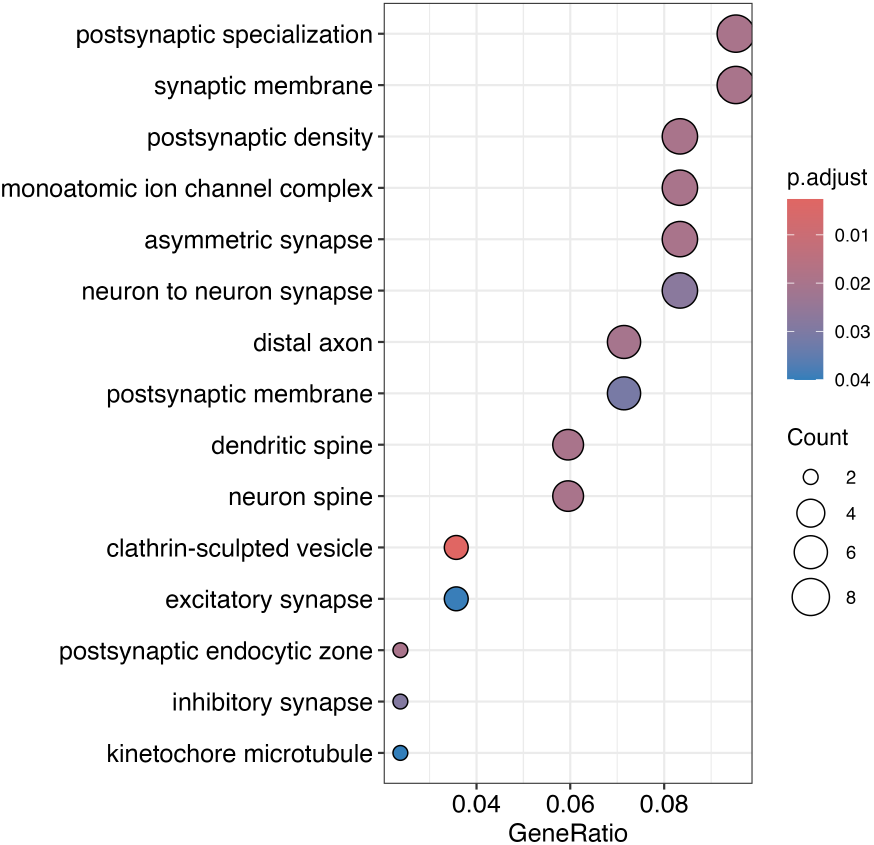
Neuronal cellular components enriched in cluster 12. Dot plot showing GO cellular component enrichment based on the top 100 genes positively defining cluster 12.

**Supplementary Fig. 8.**
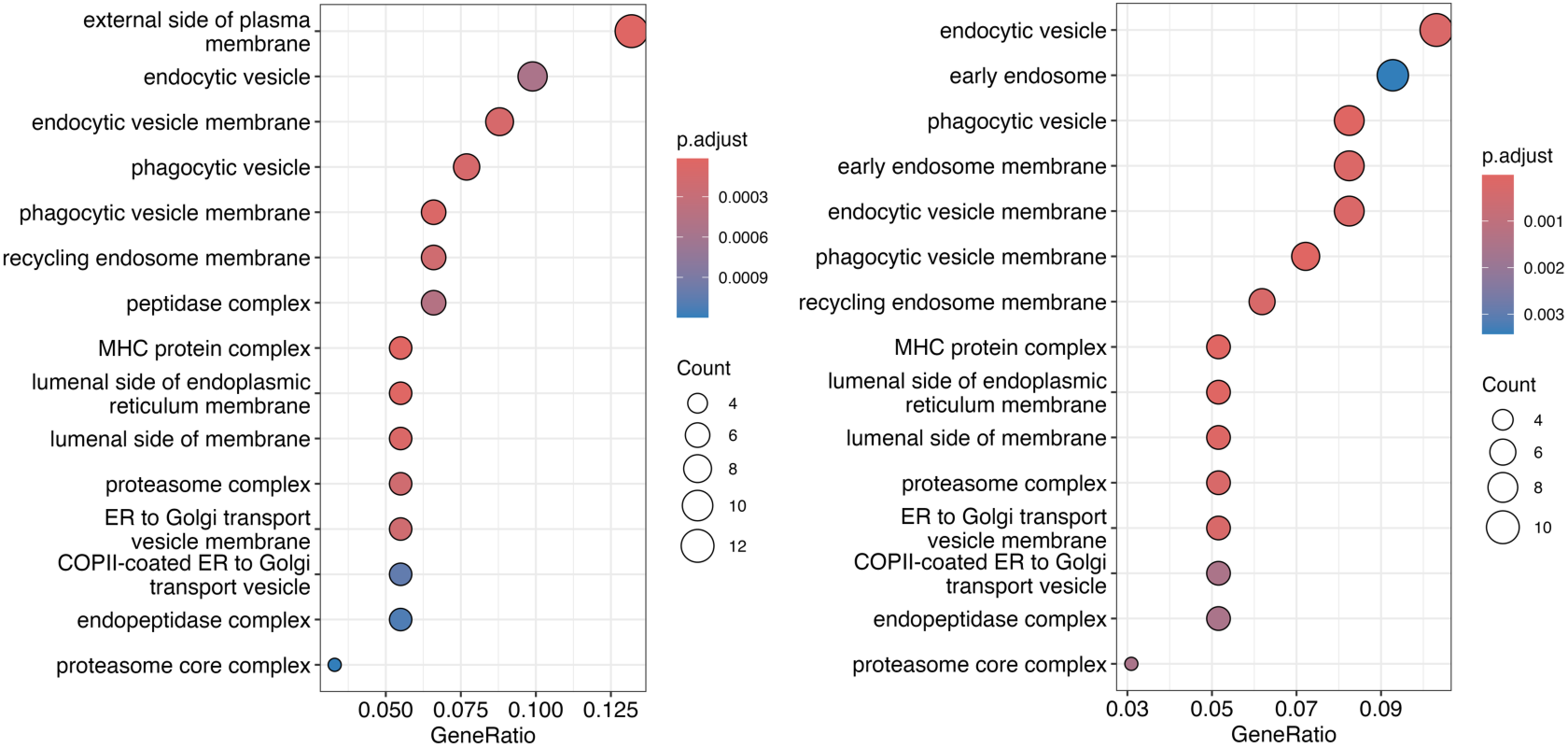
Immune-related cellular components enriched in high-dose TNF-⍺-specific RGL-like and IPC-like clusters. Dot plot showing GO cellular component enrichment based on the top 100 genes upregulated in cluster 6 as compared to cluster 5 (left). Dot plot showing GO cellular component enrichment based on the top 100 genes upregulated genes in cluster 10 as compared to cluster 7, 8, and 9 (right).

**Supplementary Fig. 9.**
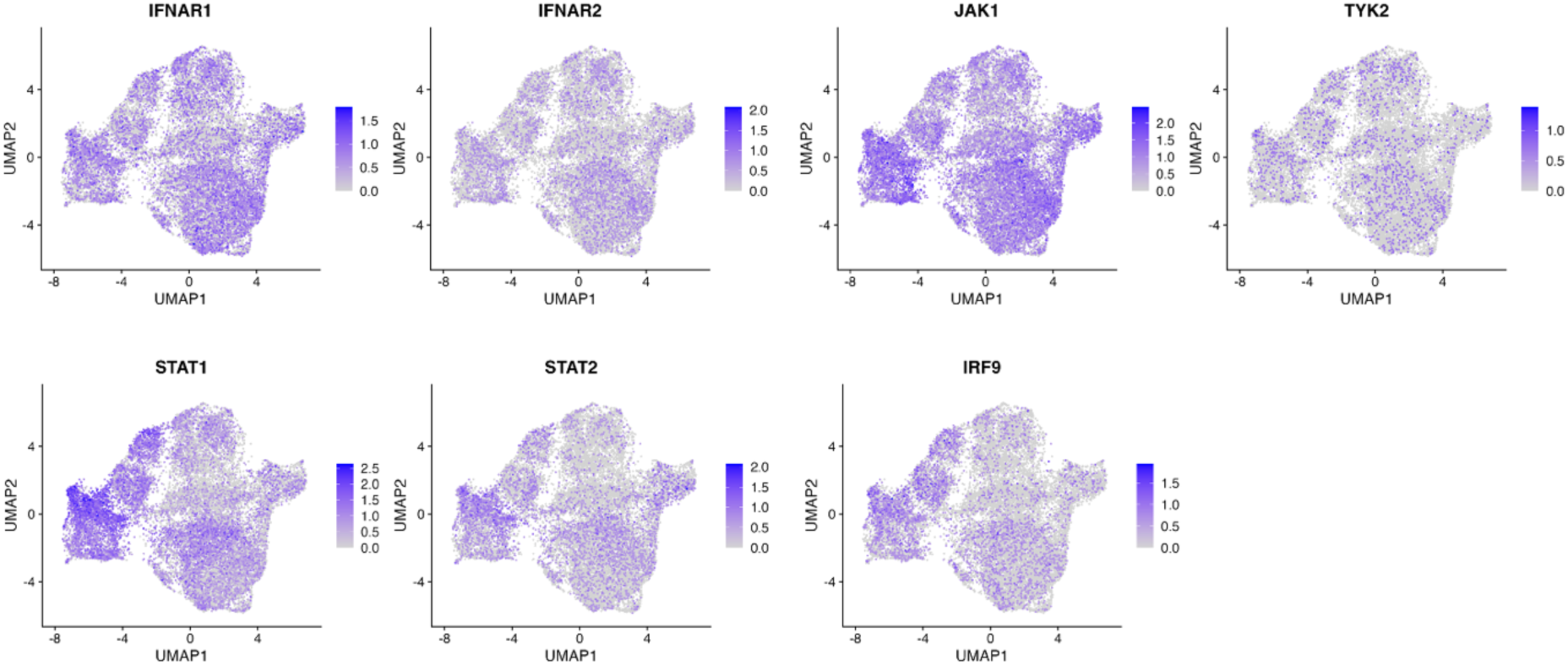
Molecular machinery for type I interferon signalling. UMAP plots showing the expression of key genes required for type I interferon signalling: *IFNAR1*, *IFNAR2*, *JAK1*, *TYK2*, *STAT1*, *STAT2*, and *IRF9*.

**Supplementary Fig. 10.**
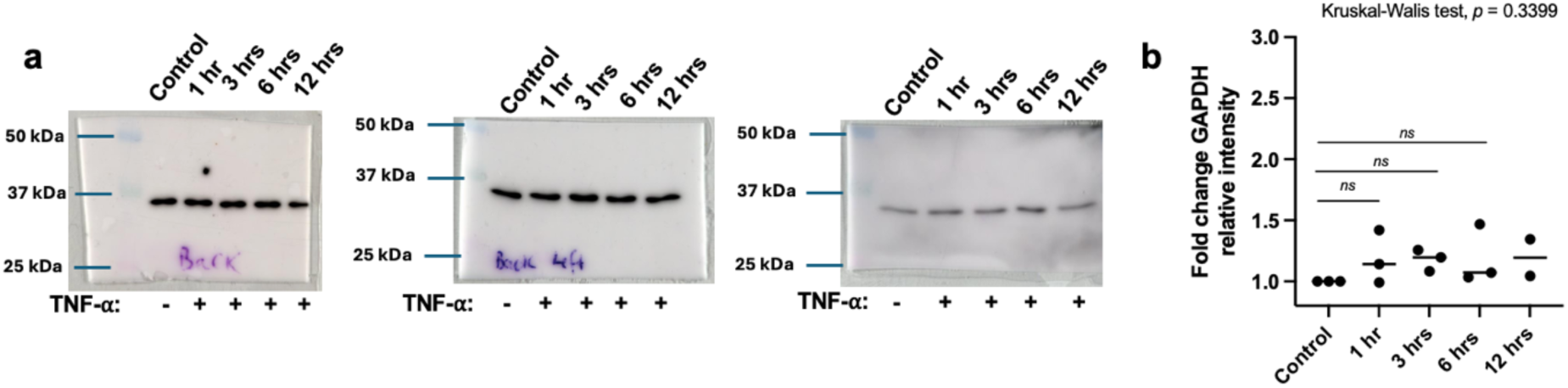
TNF-⍺ does not regulate GAPDH expression. **a,** Uncropped Western blot images for the three independent experiments probing for GAPDH. The HPCs were treated with 1 ng/ml TNF-⍺ for 1, 3, 6, or 12 hours. Molecular weight markers (Precision Plus Protein Kaleidoscope Prestained Protein Standards) are shown for reference. **b,** Quantification of GAPDH band intensities in **a**. Data represent mean ± SEM. Statistical analysis: Kruskal-Wallis test followed by Dunn’s post hoc correction.

**Supplementary Fig. 11.**
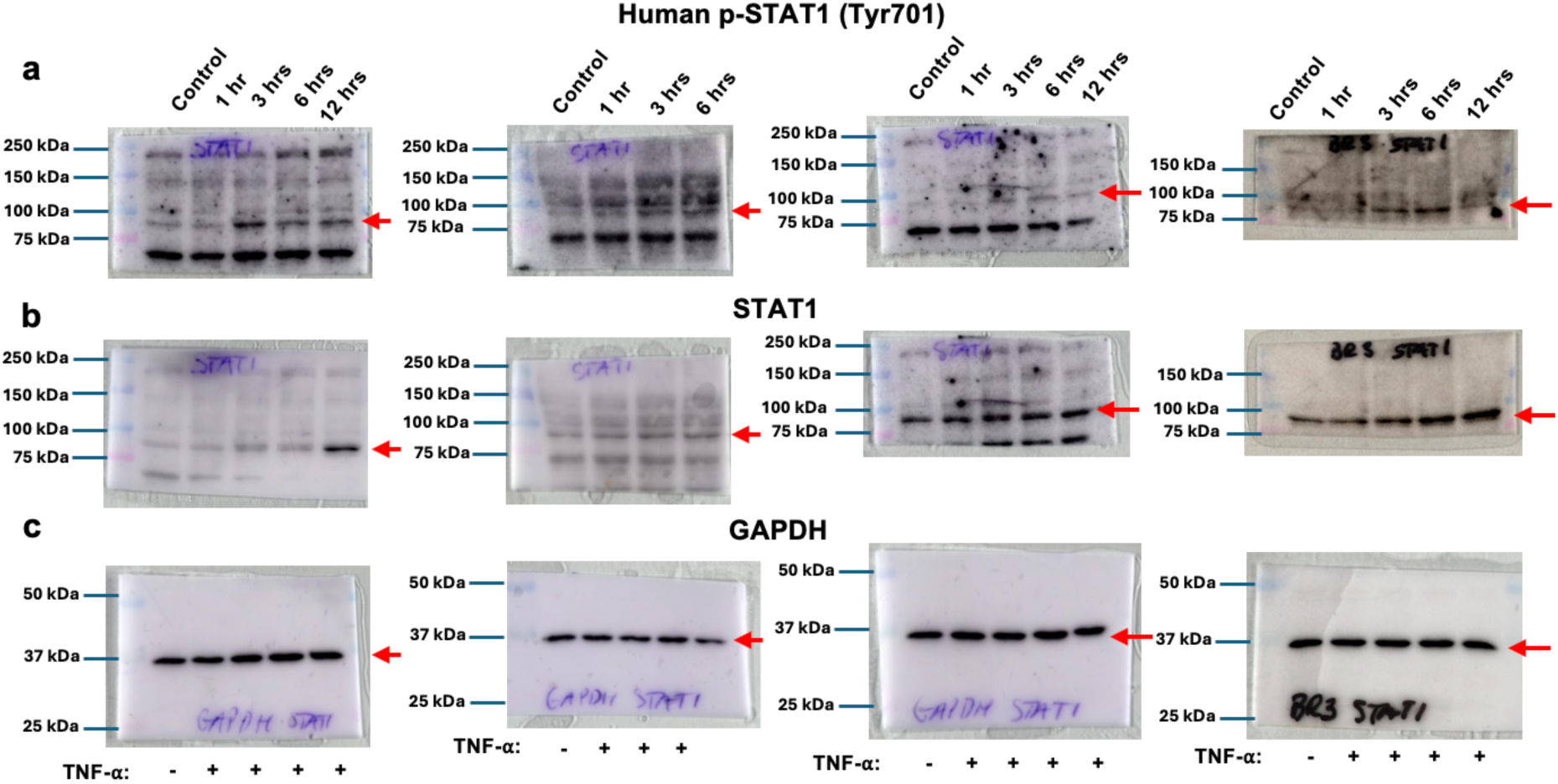
Uncropped Western blots corresponding to. Fig. 4a**-b****. a,** Uncropped Western blot images from four independent experiments probing for pSTAT1 (Tyr701). **b,** Uncropped Western blot images from four independent experiments probing for total STAT1. **c,** Uncropped Western blot images from four independent experiments probing for loading control GAPDH. HPCs were treated with 1 ng/ml TNF-⍺ for 1, 3, 6, or 12 hours. Molecular weight markers (Precision Plus Protein Kaleidoscope Prestained Protein Standards) are shown for reference. Red arrows indicate the bands of interest.

**Supplementary Fig. 12.**
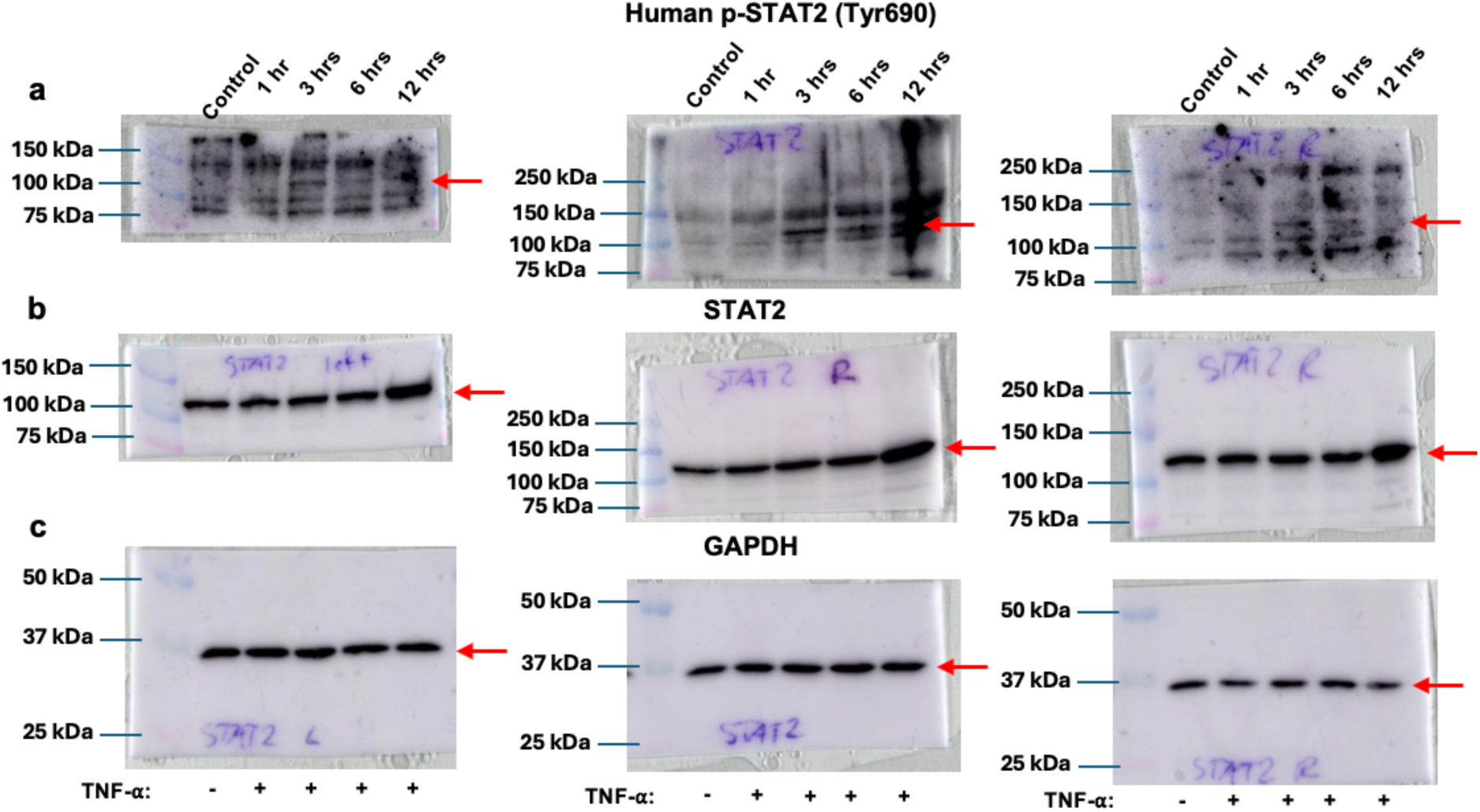
Uncropped Western blots corresponding to Extended data Fig. 1a-b. **a,** Uncropped Western blot images from four independent experiments probing for pSTAT2 (Tyr690). **b,** Uncropped Western blot images from four independent experiments probing for total STAT2. **c,** Uncropped Western blot images from four independent experiments probing for loading control GAPDH. HPCs were treated with 1 ng/ml TNF-⍺ for 1, 3, 6, or 12 hours. Molecular weight markers (Precision Plus Protein Kaleidoscope Prestained Protein Standards) are shown for reference. Red arrows indicate the band of interest.

**Supplementary Fig. 13.**
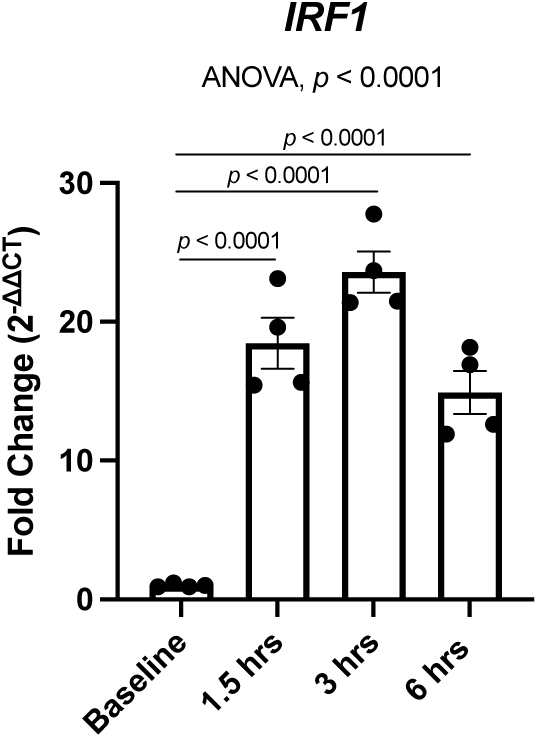
HPCs upregulate transcription factor *IRF1* in response to TNF-⍺. RT-qPCR data showing relative *IRF1* gene expression in HPCs treated ± 1 ng/ml TNF-⍺ for 1.5, 3, or 6 hours. Data is presented as mean ± SEM from four independent experiments. Statistical analysis: one-way ANOVA followed by Bonferroni’s multiple comparisons test.

**Supplementary Fig. 14.**
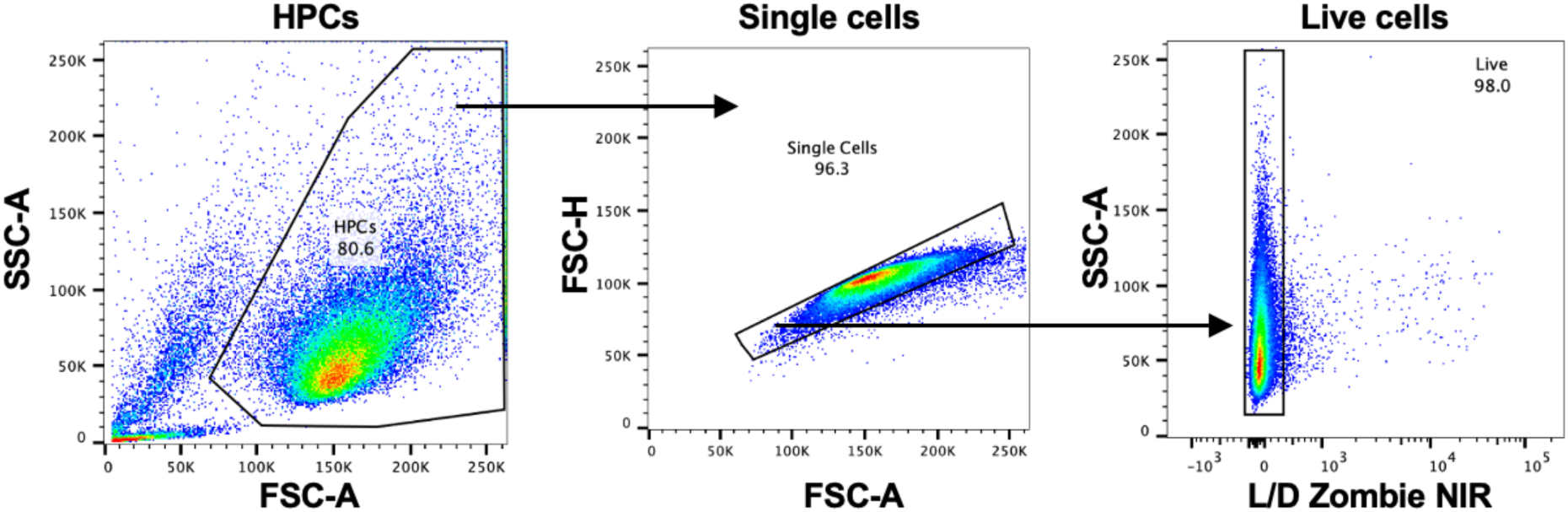
Gating strategy for assessing cell surface expression on HPCs. Representative gating strategy for analysing cell surface protein expression on live, single HPCs. Cells were first gated based on forward scatter (FSC) and side scatter (SSC) to exclude debris and identify the main population. Singlets were selected using FSC-A versus FSC-H to exclude doublets. Live cells were identified by negative selection using the Zombie NIR viability dye.

**Supplementary Fig. 15.**
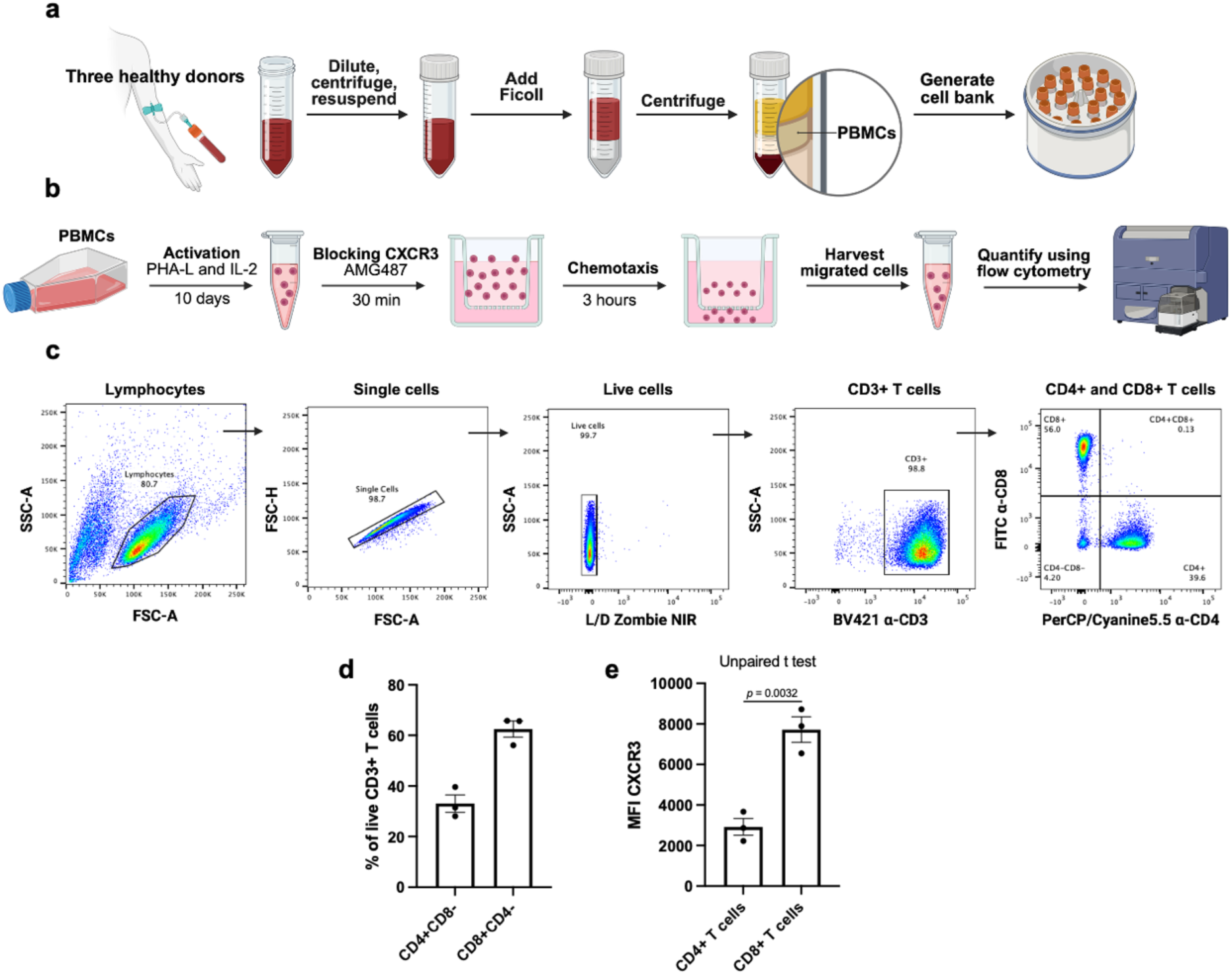
Transwell chemotaxis assay using primary human T cells to study CXCR3-dependent migration. **a,** PBMS were isolated from leukocytes cones from three healthy donors using a Ficoll-Paque density gradient, and a cell bank was generated for each donor. **b,** Schematic overview of the T cell chemotaxis assay. **c,** Representative gating strategy to identify CD4+ and CD8+ T cells. **d,** Percentage of T cell subsets following the activation protocol. Data is presented as mean ± SEM from three healthy donors. **e,** Quantification of CXCR3 MFI (median) between CD4+ T cells and CD8+ T cells. Data is presented as mean ± SEM from three healthy donors. Statistical analysis: unpaired, two-tailed, parametric t-test.

**Supplementary Fig. 16.**
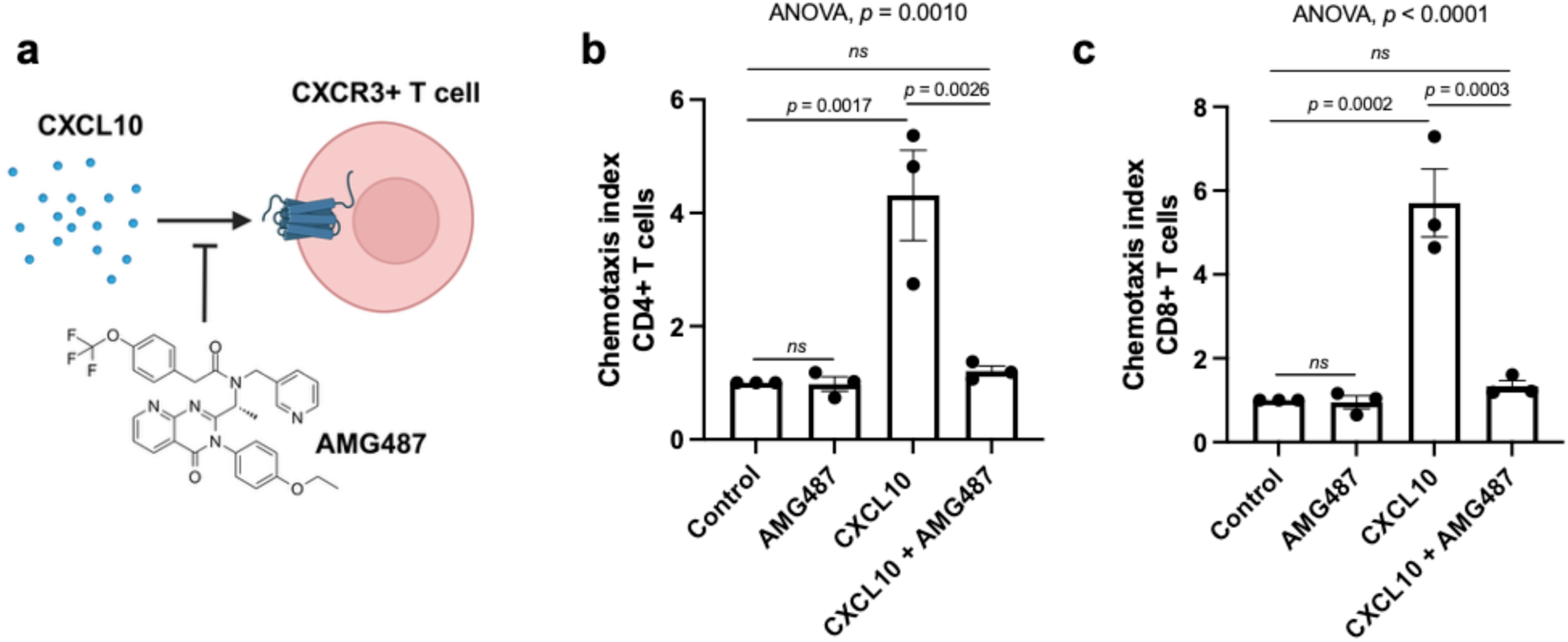
Recombinant CXCL10 promotes CXCR3-dependent T cell recruitment. **a**, CXCL10 drives T cell chemotaxis by activating CXCR3. AMG487 is a selective CXCR3 antagonist. **b**, and **c**, Quantification of the chemotactic response (chemotaxis index) of **b** CD4+ T cells and **c** CD8+ T cells treated ± 1 μM AMG487 or DMSO in response to control media or media containing 100 ng/ml recombinant hCXCL10. The chemotactic index is the ratio of the number of cells migrated in response to a stimulus, as compared with controls. Data is represented as mean ± SEM of three healthy donors. Statistical analysis: one-way ANOVA followed by Bonferroni’s multiple comparisons test.

**Supplementary Fig. 17.**
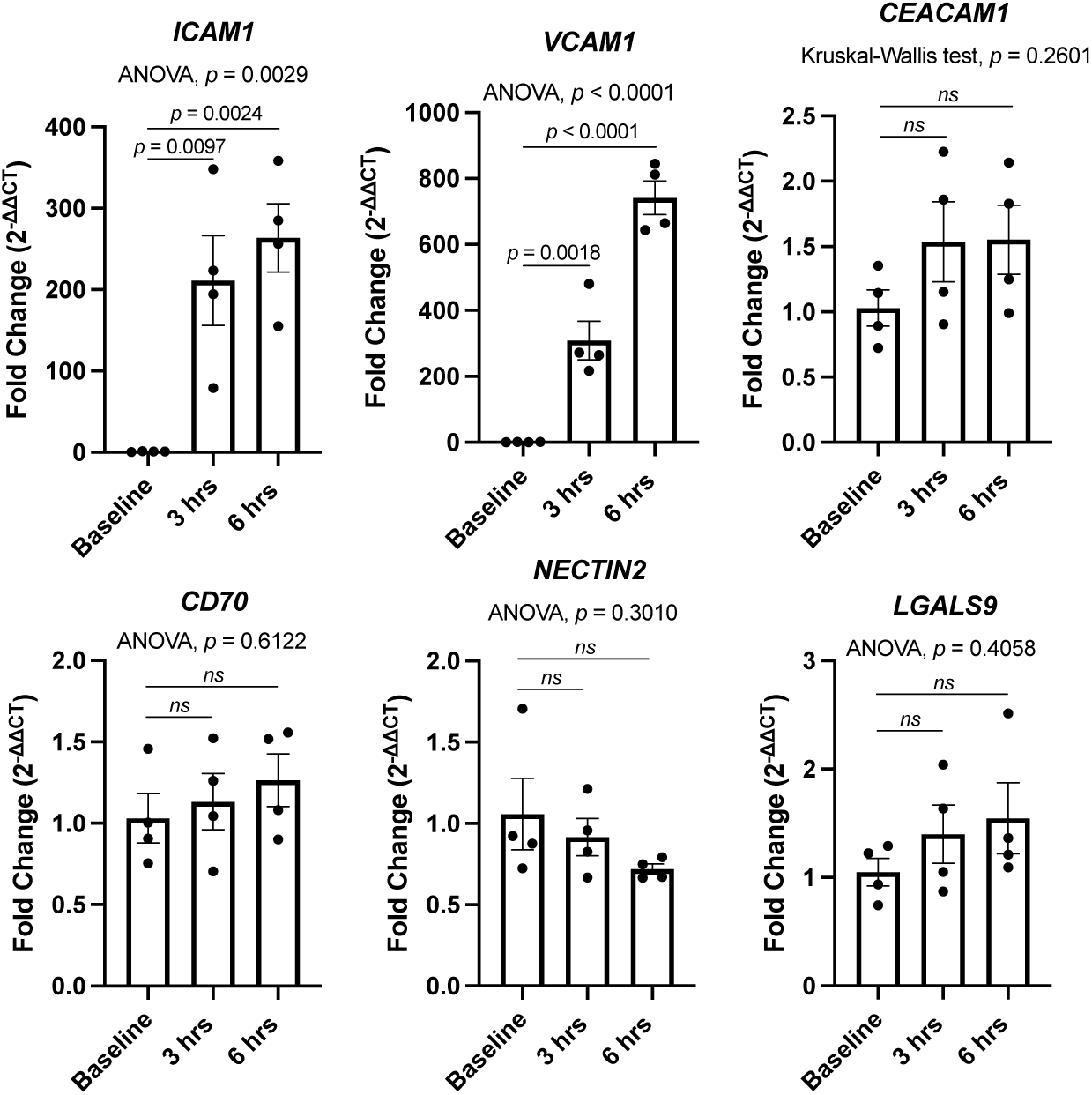
HPCs upregulate molecules involved in T cell crosstalk in response to TNF-⍺. RT-qPCR data showing relative gene expression of *ICAM1, VCAM1, CEACAM1, CD70, NECTIN2, and LGALS9* in HPCs treated ± 1 ng/ml TNF-⍺ for 1.5, 3, or 6 hours. Data is presented as mean ± SEM of four independent experiments. Statistical analysis: one-way ANOVA followed by Bonferroni’s multiple comparisons test or Kruskal-Wallis test followed by Dunn’s post hoc correction as indicated.

**Supplementary Fig. 18.**
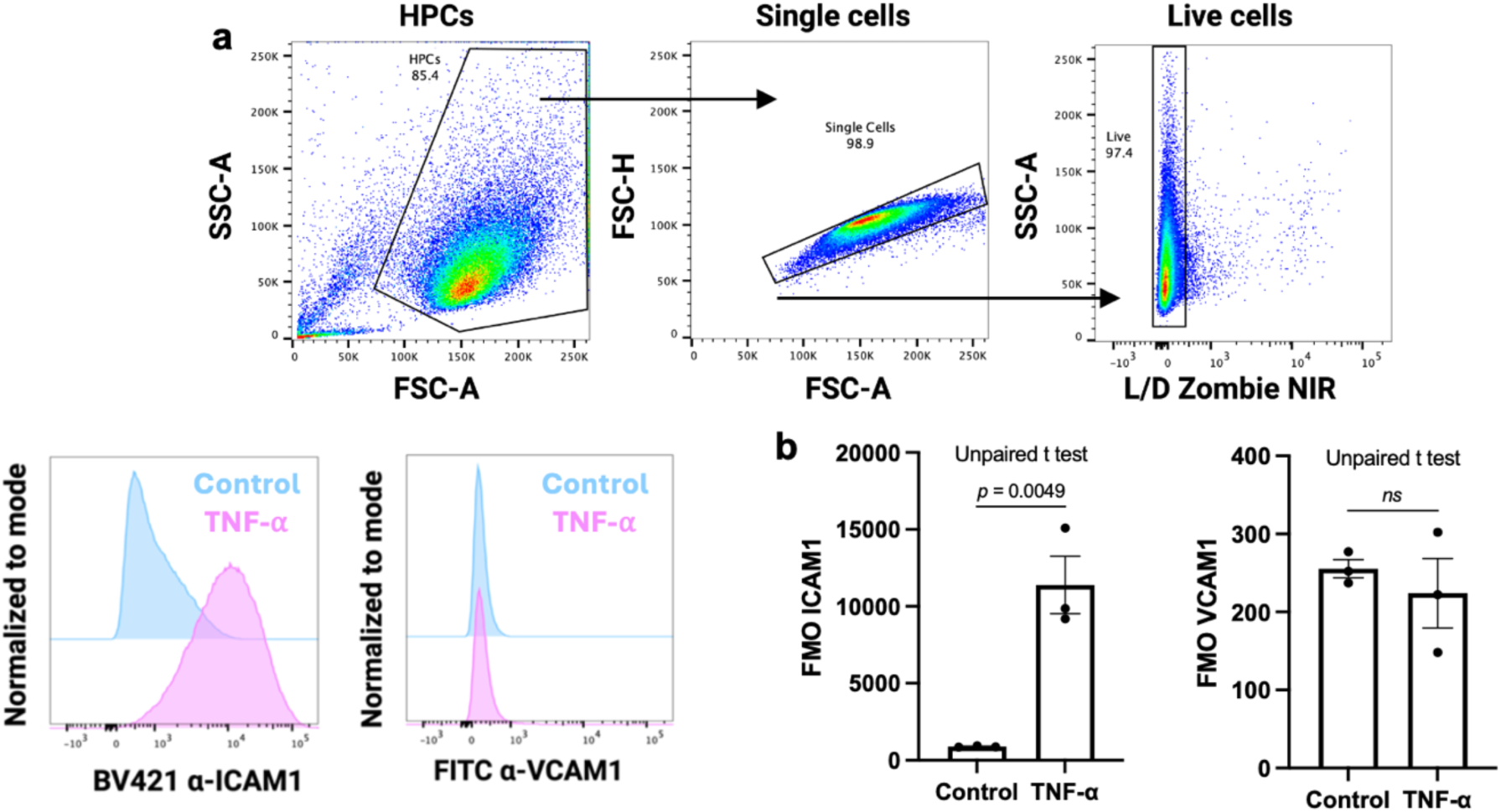
Cell surface expression of ICAM1 and VCAM1 on HPCs in response to TNF-⍺. **a,** Gating strategy for assessing cell surface protein expression on live, single HPCs. Cells were first gated by forward scatter (FSC) and side scatter (SSC) to exclude debris and identify the main population. Singlets were selected using FSC-A versus FSC-H, and live cells were identified by negative selection using the Zombie NIR viability dye. Representative histograms show ICAM1 and VCAM1 on the control-treated (blue) and TNF-⍺-treated (pink) HPCs. **b**, Quantification of ICAM1 and VCAM1 surface expression by mean fluorescence intensity (MFI) on HPCs treated ± 1 ng/ml TNF-⍺ for 24 h, measured by flow cytometry. Data is presented as mean ± SEM. Statistical analysis: two-tailed, unpaired t test.

**Supplementary Fig. 19.**
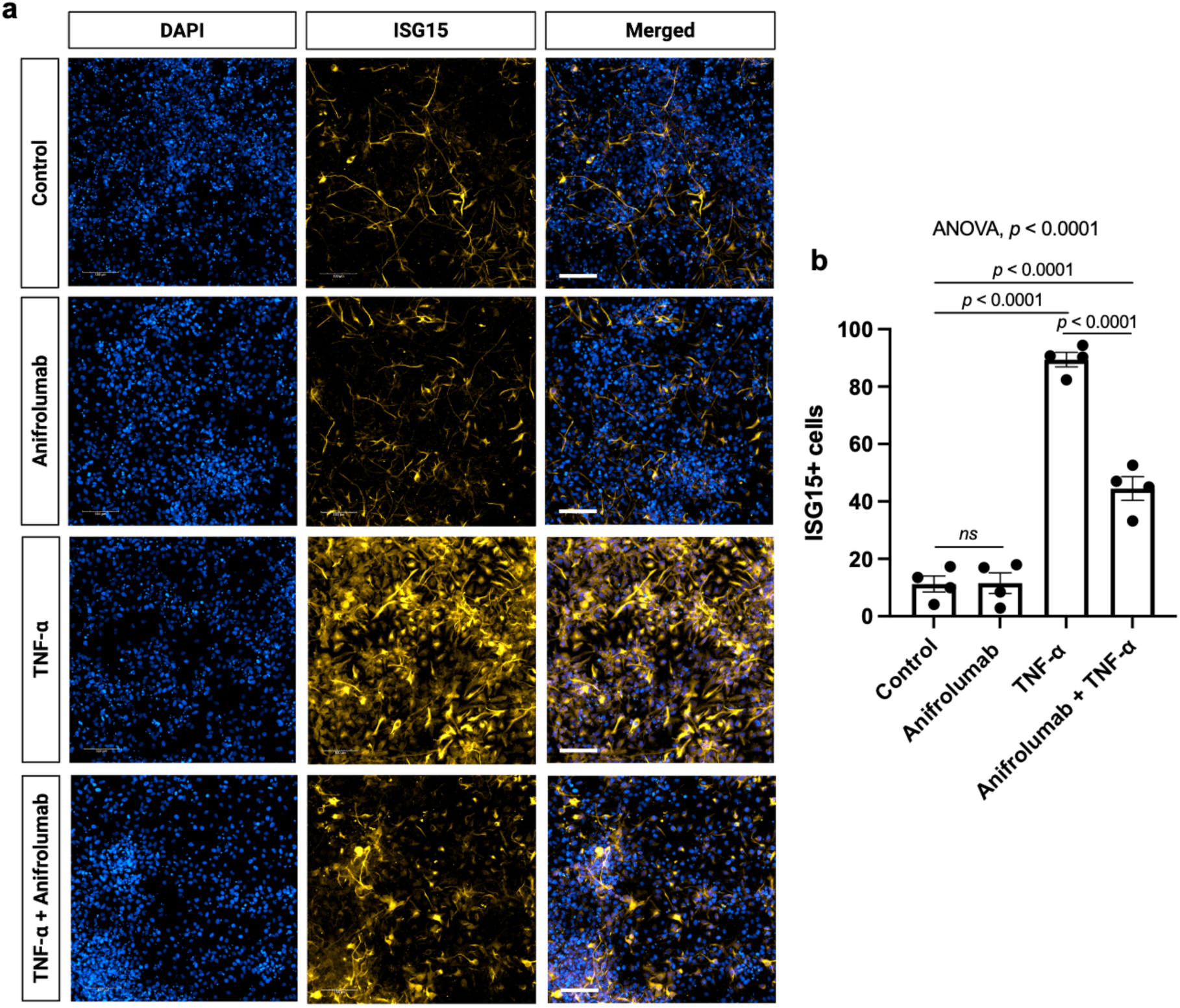
Blocking IFNAR decreases the upregulation of ISG15+ differentiating HPCs in response to chronic TNF-⍺. **a,** Representative images showing the expression of ISG15 (orange) in HPCs differentiated for seven days treated chronically with ± 1 ng/ml TNF-⍺ ± 10 µg/ml anifrolumab. Scale bar, 100 µm. **b,** Quantification of the percentage of ISG15+ cells based on panel **a**. Data represent mean ± SEM from four independent experiments. Statistical analysis: one-way ANOVA followed by Bonferroni’s multiple comparisons test.

**Supplementary Table 1.** Primary and secondary antibodies used for immunocytochemistry.

| Antibody | Manufacturer | Catalogue number | Dilution |
| --- | --- | --- | --- |
| Mouse Nestin | Chemicon | Mab5326 | 1:1000 |
| Mouse MAP2 | Abcam | Ab11267 | 1:500 |
| Rabbit DCX | Abcam | Ab18723 | 1:500 |
| Mouse STAT1 | Cell Signalling Technology | 9176 | 1:1000 |
| Rabbit ISG15 | Proteintech | 15981-1-AP | 1:500 |
| Mouse NF-κB p65 | Santa Cruz | Sc-8008 | 1:500 |
| Donkey anti-rabbit Alexa fluor 555 | Invitrogen | A-31571 | 1:500 |
| Donkey anti-rabbit Alexa fluor 488 | Invitrogen | A-21202 | 1:500 |

**Supplementary Table 2.** Thermal cycler program employed for cDNA synthesis.

| Step | Temperature (°C) | Time (minutes) |
| --- | --- | --- |
| 1 | 25 | 10 |
| 2 | 37 | 120 |
| 3 | 85 | 5 |
| 4 | 4 | ∞ |

**Supplementary Table 3.** Thermal cycling conditions employed for RT-qPCR.

| Step | Temperature (°C) | Time |
| --- | --- | --- |
| 1 | 50 | 2 minutes |
| 2 | 95 | 10 minutes |
| 3 (40 cycles) | 95 | 15 seconds |
| 4 (40 cycles) | 60 | 1 minute |
| 5 | 95 | 15 seconds |

**Supplementary Table 4.**
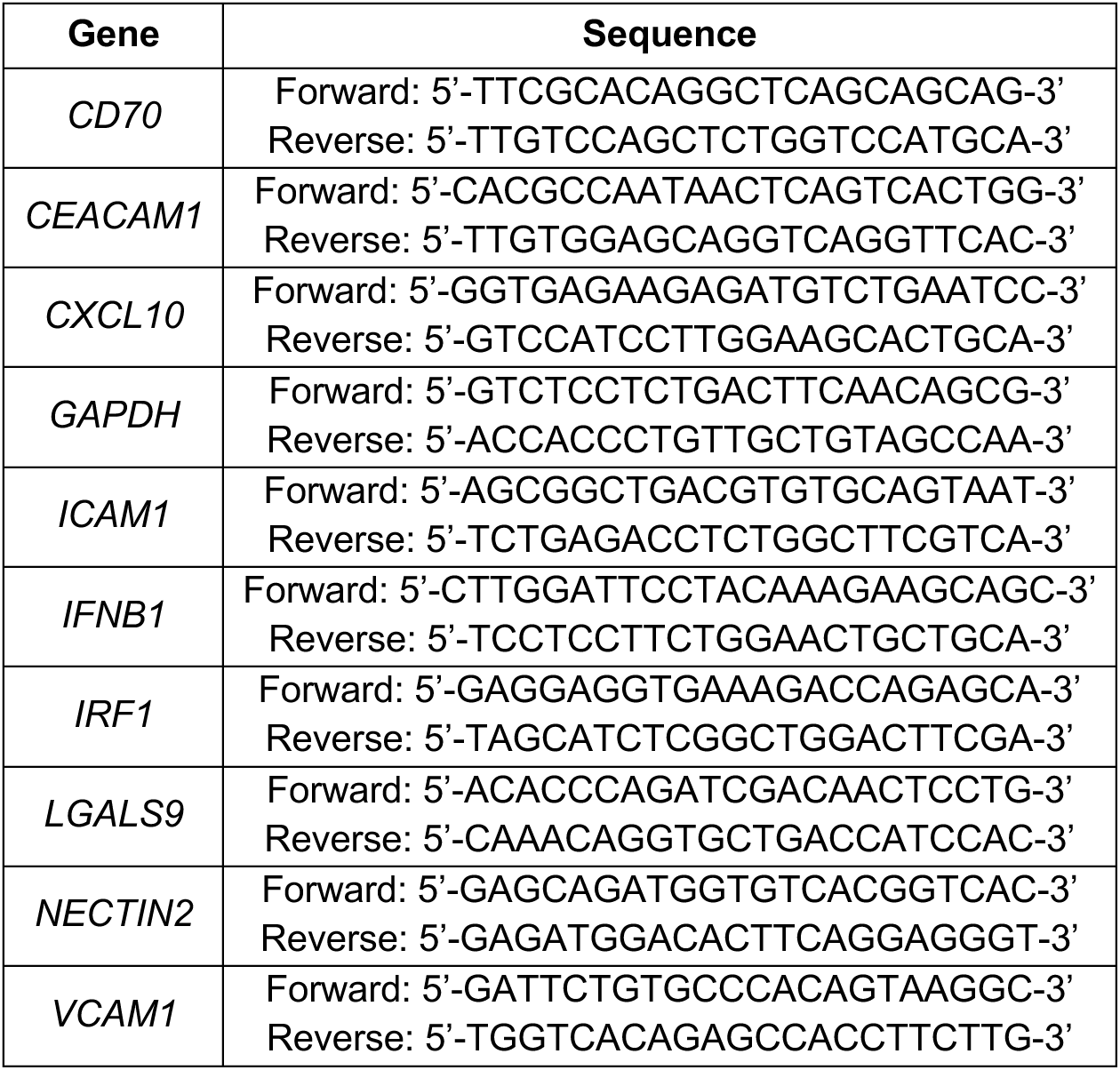
Primers used for RT-qPCR.

**Supplementary Table 5.** Antibodies used for western blot experiments.

| Target | Company | Catalogue number | Species | Dilution |
| --- | --- | --- | --- | --- |
| p-STAT1 | Cell Signalling Technology | 91715 | Rabbit | 1:1000 |
| p-STAT2 | Cell Signalling Technology | 88410 | Rabbit | 1:1000 |
| GAPDH | Proteintech | 60004-1-1g | Mouse | 1:10000 |
| STAT1 | Cell Signalling Technology | 9176 | Mouse | 1:1000 |
| STAT2 | Cell Signalling Technology | 71604 | Rabbit | 1:1000 |
| Anti-rabbit | GE | NA9340V | Donkey | 1:3000 |
| Anti-mouse | Santa Cruz Biotechnology | D1321 | Goat | 1:9000 |

**Supplementary Table 6.** Chemokines included in the human pro-inflammatory chemokine panel.

| Chemokine | Full name | Alias |
| --- | --- | --- |
| CCL2 | Chemokine (CC motif) ligand 2 | MCP-1 |
| CCL3 | Chemokine (CC motif) ligand 3 | MIP-1α |
| CCL4 | Chemokine (CC motif) ligand 4 | MIP-1β |
| CXCL1 | Chemokine (CXC motif) ligand 1 | GROα |
| CXCL5 | Chemokine (CXC motif) ligand 5 | ENA-78 |
| CXCL8 | Chemokine (CXC motif) ligand 8 | IL-8 |
| CXCL9 | Chemokine (CXC motif) ligand 9 | MIG |
| CXCL10 | Chemokine (CXC motif) ligand 10 | IP-10 |
| CXCL11 | Chemokine (CXC motif) ligand 11 | I-TAC |
| CCL11 | Chemokine (CC motif) ligand 11 | Eotaxin |
| CCL17 | Chemokine (CC motif) ligand 17 | TARC |
| CCL20 | Chemokine (CC motif) ligand 20 | MIP-3α |

**Supplementary Table 7.** Antibodies used to analyse cell surface protein expression on HPCs.

| Protein | Fluorophore | Company | Catalogue number | Dilution |
| --- | --- | --- | --- | --- |
| ICAM1 | BV421 | Biolegend | 353132 | 1:100 |
| VCAM1 | FITC | R&D | BBA22 | 1:100 |
| Tetherin (BST2) | APC | Biolegend | 348410 | 1:100 |

**Supplementary Table 8.** Antibodies used to analyse cell surface protein expression on immune cells.

| **Protein** | **Fluorophore** | **Company** | **Catalogue number** | **Dilution** |
| --- | --- | --- | --- | --- |
| CXCR3 | PE | Biolegend | 353706 | 1:100 |
| CD3 | BV421 | Biolegend | 317344 | 1:100 |
| CD4 | PerCP/Cyanine5.5 | Biolegend | 317428 | 1:100 |
| CD8 | FITC | Biolegend | 980908 | 1:100 |

## References

1. Ming, G. & Song, H. Adult Neurogenesis in the Mammalian Brain: Significant Answers and Significant Questions. Neuron 70, 687–702 (2011).

2. Altman, J. Autoradiographic investigation of cell proliferation in the brains of rats and cats. Anat Rec 145, 573–591 (1963).

3. Spalding, K. L. et al. Dynamics of Hippocampal Neurogenesis in Adult Humans. Cell 153, 1219–1227 (2013).

4. Boldrini, M. et al. Human Hippocampal Neurogenesis Persists throughout Aging. Cell Stem Cell 22, 589–599.e5 (2018).

5. Knoth, R. et al. Murine features of neurogenesis in the human hippocampus across the lifespan from 0 to 100 years. PLoS One 5, e8809 (2010).

6. Moreno-Jiménez, E. P. et al. Adult hippocampal neurogenesis is abundant in neurologically healthy subjects and drops sharply in patients with Alzheimer’s disease. Nat Med 25, 554–560 (2019).

7. Tobin, M. K. et al. Human Hippocampal Neurogenesis Persists in Aged Adults and Alzheimer’s Disease Patients. Cell Stem Cell 24, 974–982.e3 (2019).

8. Palmer, T. D. et al. Progenitor cells from human brain after death. Nature 411, 42–43 (2001).

9. Zhou, Y. et al. Molecular landscapes of human hippocampal immature neurons across lifespan. Nature 607, 527–533 (2022).

10. Dumitru, I. et al. Identification of proliferating neural progenitors in the adult human hippocampus. Science 389, 58–63 (2025).

11. Shors, T. J. et al. Neurogenesis in the adult is involved in the formation of trace memories. Nature 410, 372–376 (2001).

12. Freund, J. et al. Emergence of Individuality in Genetically Identical Mice. Science 340, 756–759 (2013).

13. Alonso, M., Petit, A.-C. & Lledo, P.-M. The impact of adult neurogenesis on affective functions: of mice and men. Mol Psychiatry 29, 2527–2542 (2024).

14. Babcock, K. R., Page, J. S., Fallon, J. R. & Webb, A. E. Adult Hippocampal Neurogenesis in Aging and Alzheimer’s Disease. Stem Cell Reports 16, 681–693 (2021).

15. Berger, T., Lee, H., Young, A. H., Aarsland, D. & Thuret, S. Adult Hippocampal Neurogenesis in Major Depressive Disorder and Alzheimer’s Disease. Trends Mol Med 26, 803–818 (2020).

16. Hill, A. S., Sahay, A. & Hen, R. Increasing Adult Hippocampal Neurogenesis is Sufficient to Reduce Anxiety and Depression-Like Behaviors. Neuropsychopharmacology 40, 2368–2378 (2015).

17. Li, Y.-D. et al. Activation of hypothalamic-enhanced adult-born neurons restores cognitive and affective function in Alzheimer’s disease. Cell Stem Cell 30, 415–432.e6 (2023).

18. Heneka, M. T. et al. Neuroinflammation in Alzheimer disease. Nat Rev Immunol 25, 321–352 (2025).

19. Zhang, W., Sun, H.-S., Wang, X., Dumont, A. S. & Liu, Q. Cellular senescence, DNA damage, and neuroinflammation in the aging brain. Trends Neurosci 47, 461–474 (2024).

20. Beurel, E., Toups, M. & Nemeroff, C. B. The Bidirectional Relationship of Depression and Inflammation: Double Trouble. Neuron 107, 234–256 (2020).

21. Swardfager, W. et al. A Meta-Analysis of Cytokines in Alzheimer’s Disease. Biol Psychiatry 68, 930–941 (2010).

22. Holmes, C. et al. Systemic inflammation and disease progression in Alzheimer disease. Neurology 73, 768–774 (2009).

23. Brymer, K. J., Romay-Tallon, R., Allen, J., Caruncho, H. J. & Kalynchuk, L. E. Exploring the Potential Antidepressant Mechanisms of TNFα Antagonists. Front Neurosci 13, (2019).

24. Enache, D., Pariante, C. M. & Mondelli, V. Markers of central inflammation in major depressive disorder: A systematic review and meta-analysis of studies examining cerebrospinal fluid, positron emission tomography and post-mortem brain tissue. Brain Behav Immun 81, 24–40 (2019).

25. Borsini, A., Zunszain, P. A., Thuret, S. & Pariante, C. M. The role of inflammatory cytokines as key modulators of neurogenesis. Trends Neurosci 38, 145–57 (2015).

26. Monje, M. L., Toda, H. & Palmer, T. D. Inflammatory Blockade Restores Adult Hippocampal Neurogenesis. Science 302, 1760–1765 (2003).

27. Chen, M., Reed, R. R. & Lane, A. P. Chronic Inflammation Directs an Olfactory Stem Cell Functional Switch from Neuroregeneration to Immune Defense. Cell Stem Cell 25, 501–513.e5 (2019).

28. Dulken, B. W. et al. Single-cell analysis reveals T cell infiltration in old neurogenic niches. Nature 571, 205–210 (2019).

29. Wu, Y. et al. Multimodal transcriptomics reveal neurogenic aging trajectories and age-related regional inflammation in the dentate gyrus. Nat Neurosci 28, 415–430 (2025).

30. Laurent, C. et al. Hippocampal T cell infiltration promotes neuroinflammation and cognitive decline in a mouse model of tauopathy. Brain 140, 184–200 (2017).

31. Walgrave, H. et al. Restoring miR-132 expression rescues adult hippocampal neurogenesis and memory deficits in Alzheimer’s disease. Cell Stem Cell 28, 1805–1821.e8 (2021).

32. Borsini, A. et al. Neurogenesis is disrupted in human hippocampal progenitor cells upon exposure to serum samples from hospitalized COVID-19 patients with neurological symptoms. Mol Psychiatry 27, 5049–5061 (2022).

33. Du Preez, A. et al. The serum metabolome mediates the concert of diet, exercise, and neurogenesis, determining the risk for cognitive decline and dementia. Alzheimer’s & Dementia 18, 654–675 (2022).

34. Maruszak, A. et al. Predicting progression to Alzheimer’s disease with human hippocampal progenitors exposed to serum. Brain 146, 2045–2058 (2023).

35. Anacker, C. et al. Role for the kinase SGK1 in stress, depression, and glucocorticoid effects on hippocampal neurogenesis. Proceedings of the National Academy of Sciences 110, 8708–8713 (2013).

36. de Lucia, C. et al. Lifestyle mediates the role of nutrient-sensing pathways in cognitive aging: cellular and epidemiological evidence. Commun Biol 3, 157 (2020).

37. Gutierrez, H. & Davies, A. M. Regulation of neural process growth, elaboration and structural plasticity by NF-κB. Trends Neurosci 34, 316–325 (2011).

38. Carvajal Ibañez, D., et al. Interferon regulates neural stem cell function at all ages by orchestrating mTOR and cell cycle. EMBO Mol Med 15, (2023).

39. Borsini, A. et al. Interferon-Alpha Reduces Human Hippocampal Neurogenesis and Increases Apoptosis via Activation of Distinct STAT1-Dependent Mechanisms. Int J Neuropsychopharmacol 21, 187–200 (2018).

40. Yarilina, A., Park-Min, K.-H., Antoniv, T., Hu, X. & Ivashkiv, L. B. TNF activates an IRF1-dependent autocrine loop leading to sustained expression of chemokines and STAT1-dependent type I interferon–response genes. Nat Immunol 9, 378–387 (2008).

41. Venkatesh, D. et al. Endothelial TNF Receptor 2 Induces IRF1 Transcription Factor-Dependent Interferon-β Autocrine Signaling to Promote Monocyte Recruitment. Immunity 38, 1025–1037 (2013).

42. Bonelli, M. et al. IRF1 is critical for the TNF-driven interferon response in rheumatoid fibroblast-like synoviocytes. Exp Mol Med 51, 1–11 (2019).

43. Riggs, J. M. et al. Characterisation of anifrolumab, a fully human anti-interferon receptor antagonist antibody for the treatment of systemic lupus erythematosus. Lupus Sci Med 5, e000261 (2018).

44. Thompson, J. E. et al. Photochemical preparation of a pyridone containing tetracycle: A jak protein kinase inhibitor. Bioorg Med Chem Lett 12, 1219–1223 (2002).

45. Chauquet, S. et al. Exercise rejuvenates microglia and reverses T cell accumulation in the aged female mouse brain. Aging Cell 23 (2024).

46. Krukowski, K. et al. Small molecule cognitive enhancer reverses age-related memory decline in mice. Elife 9, (2020).

47. Unger, M. S. et al. CD8+ T-cells infiltrate Alzheimer’s disease brains and regulate neuronal- and synapse-related gene expression in APP-PS1 transgenic mice. Brain Behav Immun 89, 67–86 (2020).

48. Gate, D. et al. Clonally expanded CD8 T cells patrol the cerebrospinal fluid in Alzheimer’s disease. Nature 577, 399–404 (2020).

49. Merlini, M., Kirabali, T., Kulic, L., Nitsch, R. M. & Ferretti, M. T. Extravascular CD3+ T Cells in Brains of Alzheimer Disease Patients Correlate with Tau but Not with Amyloid Pathology: An Immunohistochemical Study. Neurodegener Dis 18, 49–56 (2018).

50. Groom, J. R. & Luster, A. D. CXCR3 ligands: redundant, collaborative and antagonistic functions. Immunol Cell Biol 89, 207–215 (2011).

51. Hsu, W.-L., Ma, Y.-L., Hsieh, D.-Y., Liu, Y.-C. & Lee, E. H. STAT1 Negatively Regulates Spatial Memory Formation and Mediates the Memory-Impairing Effect of Aβ. Neuropsychopharmacology 39, 746–758 (2014).

52. Imitola, J. et al. Stat1 is an inducible transcriptional repressor of neural stem cells self-renewal program during neuroinflammation. Front Cell Neurosci 17, (2023).

53. Zheng, L.-S. et al. Mechanisms for Interferon-α-Induced Depression and Neural Stem Cell Dysfunction. Stem Cell Reports 3, 73–84 (2014).

54. Baruch, K. et al. Aging-induced type I interferon response at the choroid plexus negatively affects brain function. Science 346, 89–93 (2014).

55. Chen, Z. & Palmer, T. D. Differential roles of TNFR1 and TNFR2 signaling in adult hippocampal neurogenesis. Brain Behav Immun 30, 45–53 (2013).

56. Liu, Y.-P., Lin, H.-I. & Tzeng, S.-F. Tumor necrosis factor-α and interleukin-18 modulate neuronal cell fate in embryonic neural progenitor culture. Brain Res 1054, 152–158 (2005).

57. Keohane, A., Ryan, S., Maloney, E., Sullivan, A. M. & Nolan, Y. M. Tumour necrosis factor-α impairs neuronal differentiation but not proliferation of hippocampal neural precursor cells: Role of Hes1. Molecular and Cellular Neuroscience 43, 127–135 (2010).

58. Neumann, H. et al. Tumor Necrosis Factor Inhibits Neurite Outgrowth and Branching of Hippocampal Neurons by a Rho-Dependent Mechanism. The Journal of Neuroscience 22, 854–862 (2002).

59. Sheng, W. S. et al. TNF-α-induced chemokine production and apoptosis in human neural precursor cells. J Leukoc Biol 78, 1233–1241 (2005).

60. McNab, F., Mayer-Barber, K., Sher, A., Wack, A. & O’Garra, A. Type I interferons in infectious disease. Nat Rev Immunol 15, 87–103 (2015).

61. Ninh, V. K. et al. Spatially clustered type I interferon responses at injury borderzones. Nature 633, 174–181 (2024).

62. Ashby, K. M., et al. Sterile production of interferons in the thymus affects T cell repertoire selection. Sci Immunol 9, (2024).

63. Tliba, O. et al. Tumor Necrosis Factor α Modulates Airway Smooth Muscle Function via the Autocrine Action of Interferon β. Journal of Biological Chemistry 278, 50615–50623 (2003).

64. Escoubas, C. C. et al. Type I interferon responsive microglia shape cortical development and behavior. Preprint at 10.1101/2021.04.29.441889 (2021).

65. de Weerd, N. A. & Nguyen, T. The interferons and their receptors—distribution and regulation. Immunol Cell Biol 90, 483–491 (2012).

66. Deczkowska, A. et al. Mef2C restrains microglial inflammatory response and is lost in brain ageing in an IFN-I-dependent manner. Nat Commun 8, 717 (2017).

67. Roy, E. R. et al. Type I interferon response drives neuroinflammation and synapse loss in Alzheimer disease. Journal of Clinical Investigation 130, 1912– 1930 (2020).

68. Roy, E. R. et al. Concerted type I interferon signaling in microglia and neural cells promotes memory impairment associated with amyloid β plaques. Immunity 55, 879–894.e6 (2022).

69. Licht-Murava, A. et al. Astrocytic TDP-43 dysregulation impairs memory by modulating antiviral pathways and interferon-inducible chemokines. Sci Adv 9, (2023).

70. Wang, Q. et al. Brain derived β-interferon is a potential player in Alzheimer’s disease pathogenesis and cognitive impairment. Alzheimers Res Ther 16, 271 (2024).

71. Hur, J.-Y. et al. The innate immunity protein IFITM3 modulates γ-secretase in Alzheimer’s disease. Nature 586, 735–740 (2020).

72. Stokes, C. et al. The human neural cell atlas of Zika virus infection in developing brain tissue. Cell Rep Med 6, 102189 (2025).

73. Fernando, N. et al. Single-cell multiomic analysis reveals the involvement of Type I interferon-responsive CD8+ T cells in amyloid beta-associated memory loss. Preprint at 10.1101/2023.03.18.533293 (2023).

74. Mason, H. D. & McGavern, D. B. How the immune system shapes neurodegenerative diseases. Trends Neurosci 45, 733–748 (2022).

75. Chen, X. & Holtzman, D. M. Emerging roles of innate and adaptive immunity in Alzheimer’s disease. Immunity 55, 2236–2254 (2022).

76. Groh, J. et al. Microglia activation orchestrates CXCL10-mediated CD8+ T cell recruitment to promote aging-related white matter degeneration. Nat Neurosci 28, 1160–1173 (2025).

77. Ma, W. et al. Type I interferon response in astrocytes promotes brain metastasis by enhancing monocytic myeloid cell recruitment. Nat Commun 14, 2632 (2023).

78. Nelson, T. E. & Gruol, D. L. The chemokine CXCL10 modulates excitatory activity and intracellular calcium signaling in cultured hippocampal neurons. J Neuroimmunol 156, 74–87 (2004).

79. Petrisko, T. J. et al. Neuronal CXCL10/CXCR3 Axis Mediates the Induction of Cerebral Hyperexcitability by Peripheral Viral Challenge. Front Neurosci 14, (2020).

80. Kalamakis, G. et al. Quiescence Modulates Stem Cell Maintenance and Regenerative Capacity in the Aging Brain. Cell 176, 1407–1419.e14 (2019).

81. Du Preez, A. et al. Impaired hippocampal neurogenesis in vitro is modulated by dietary-related endogenous factors and associated with depression in a longitudinal ageing cohort study. Mol Psychiatry 27, 3425–3440 (2022).

82. Farmand, S., Ahmed, E., Zawar, H. A. & Thuret, S. Selenium deficiency negatively affects survival and integrity of human hippocampal progenitor cells. Aging Brain 7, 100138 (2025).

83. Wu, T. et al. clusterProfiler 4.0: A universal enrichment tool for interpreting omics data. The Innovation 2, 100141 (2021).

84. Badia-I-Mompel, P. et al. decoupleR: ensemble of computational methods to infer biological activities from omics data. Bioinformatics advances 2, vbac016 (2022).

85. Müller-Dott, S. et al. Expanding the coverage of regulons from high-confidence prior knowledge for accurate estimation of transcription factor activities. Nucleic Acids Res 51, 10934–10949 (2023).

86. Türei, D. et al. Integrated intra- and intercellular signaling knowledge for multicellular omics analysis. Mol Syst Biol 17, (2021).

